# Neurobehavioral effects of focused ultrasound-mediated blood-brain barrier opening

**DOI:** 10.64898/2026.08.12.744037

**Authors:** Sophie K. Elvig, Alexandra A. Seas, Pavlos Anastasiadis, Steffen B. E. Wolff

**Author notes:** Authors contributed equally (Sophie K. Elvig, Alexandra A. Seas). Co-corresponding authors: Steffen Wolff, Ph.D., Pavlos Anastasiadis, Ph.D.

## Abstract

Microbubble-enhanced focused ultrasound (MB-FUS) enables noninvasive blood-brain barrier (BBB) opening to improve the delivery of drugs and other therapeutics to the brain, supporting more effective treatment of brain disorders. The benefits of this rapidly advancing and highly versatile technology have been demonstrated in clinical trials, sparking a growing interest in expanding FUS applications that require higher intensity treatments, larger targeted brain volumes and larger therapeutics. However, in-depth safety profiling of such treatments has not been done and is limited in the clinic. Preclinical studies have been restricted in their readouts, focusing on acute imaging and simple behaviors. To address the need for more holistic safety profiling, we present a novel preclinical workflow for determining adverse effects of MB-FUS in rats, both acutely and long-term, by combining MRI, histology, and a custom motor task which provides fine-scaled readouts for complex learned behavior. Using this approach and taking advantage of our previous delineation of the relevant circuitry, we show dose- and target-dependent adverse effects in high dosing regimens. All prescribed acoustic doses opened the BBB; but while low doses had no overt adverse effects, high doses targeting the involved circuitry had severe effects on both behavior and brain tissue integrity. These effects persisted for weeks and recovered over differing time courses, with tissue disruptions and behavioral changes outlasting general performance deficits. Our results reinforce the need for multimodal, highly sensitive, and longitudinal readouts to holistically characterize adverse effects of MB-FUS, allowing for its safe use across a wide range of applications.

**Significance:** Microbubble-enhanced focused ultrasound is emerging as a powerful noninvasive approach for delivering drugs, genes, and cell therapies through the blood-brain barrier, sparking broad interest in expanding clinical and preclinical applications. However, its effects on complex neurological function, especially for higher dosing regimens, remain poorly defined. To address this, we have established a novel multimodal preclinical strategy for defining functional safety limits and guiding clinical translation, integrating sensitive behavioral testing with MRI, histology and acoustic emissions analysis. We show dose- and target-dependent impairments in complex learned motor behavior and evidence of possible brain injury, with substantially different recovery time courses. Together, our findings demonstrate the importance of a multimodal, longitudinal approach for comprehensively characterizing treatment-related adverse effects and evaluating safety.

## INTRODUCTION

Focused ultrasound (FUS) is a rapidly evolving and highly versatile treatment technology, especially in the fields of neurology and neuro-oncology. It enables noninvasive treatment with exceptional spatial precision to target brain regions and circuits that are involved in or disrupted by clinically challenging disorders including Parkinson’s Disease, Alzheimer’s Disease, and High-Grade Glioma (1–6). FUS directs acoustic energy within a spatially well-defined, millimeter-precise focal volume, generating a mechanical perturbation of the targeted brain tissue. The resulting effects depend on treatment parameters, including frequency, duty cycle, pulse width, and pulse repetition frequency. High-intensity FUS is used for targeted tissue thermoablation, for example, in neurological tremor conditions, including Essential Tremor and Parkinson’s Disease (7–9). More recently, microbubble-enhanced FUS (MB-FUS) has dramatically expanded the versatility of FUS and holds the promise to address a critical treatment need by allowing for the transient modulation of blood-brain barrier (BBB) permeability (10) (***Fig. 1***). MBs are micron-sized gas-filled lipid vesicles which are traditionally used as contrast agents in ultrasound imaging (11). In MB-FUS, a focused beam of acoustic energy induces size oscillations in intravascularly injected MBs, which in turn disrupt the neurovascular unit and cause transient BBB opening (12). Through this mechanism, MB-FUS can enhance brain delivery of drugs and other agents that cannot cross the intact BBB, helping overcome one of the biggest obstacles to treating a wide spectrum of central nervous system indications with previously underutilized drugs. Taking advantage of this potential, the field has experienced a remarkable expansion of clinical treatment approaches over the past two decades (13).

**Figure 1.**
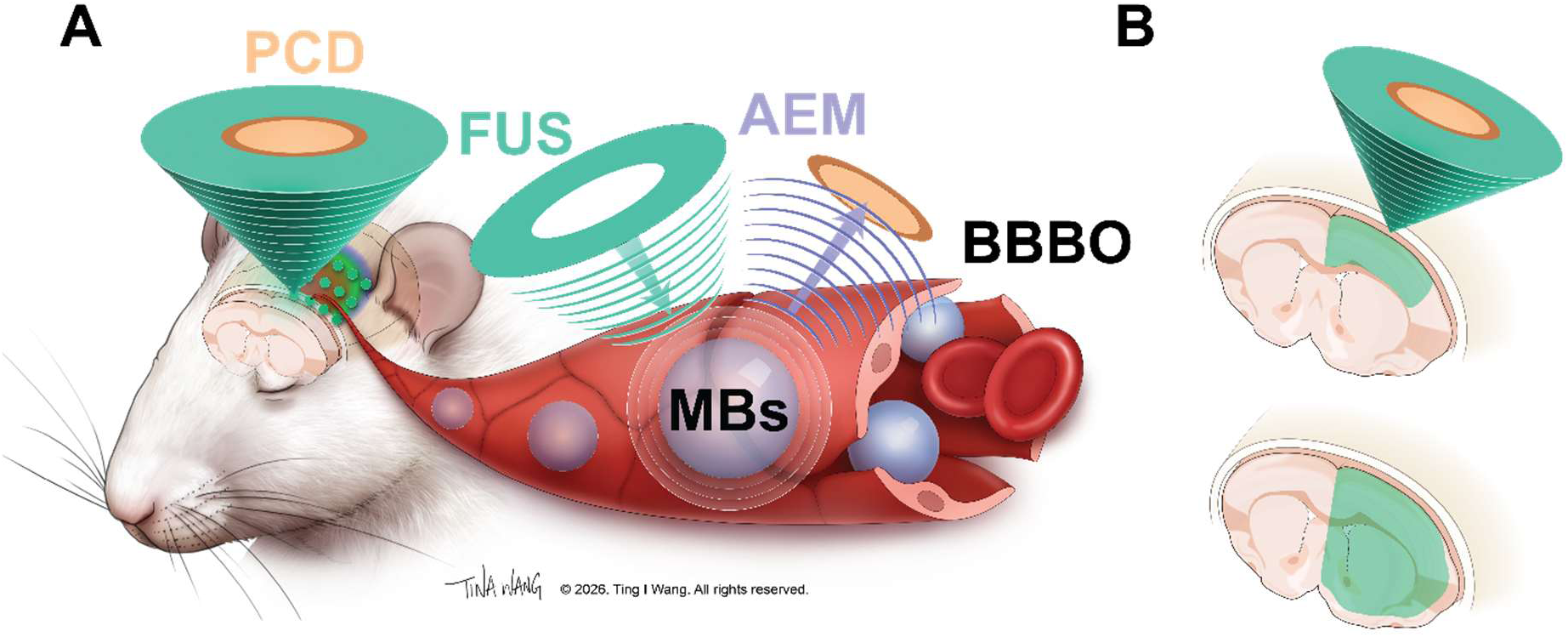
MB-FUS treatment summary. **A** Animals were treated with a 5×4 spot grid using a singleelement FUS transducer with simultaneous infusion of microbubbles (MBs). Passive cavitation data (PCD) from bubble oscillations was collected for acoustic emissions monitoring (AEM). Blood-brain barrier opening (BBBO) was confirmed radiographically with T1 – contrast enhanced magnetic resonance imaging (MRI). B Altering of the transducer bolus allowed us to target either only cortical (superficial; top) or cortical and subcortical (deep; bottom) brain areas for treatment.

A critical prerequisite for taking full advantage of MB-FUS’s potential and for establishing and expanding it as a versatile treatment option is to fully characterize its safety profile. To date, 36 early-phase clinical trials have been completed or are currently underway to assess the safety and efficacy of MB-FUS for opening the BBB and for improving drug delivery to the brain (1, 14–18) (see Methods for trial inclusion criteria). Enhancing drug delivery for newly developed, larger therapeutics beyond small-molecule drugs (e.g., gene therapy, antibodies, CAR T cells), requires higher intensity FUS settings to adequately increase permeability of the brain microvasculature. However, such a dose escalation also increases the risk of acute microvascular injury, potentially leading to microhemorrhages and neuroinflammatory activation (19, 20). Balancing BBB opening with these adverse side effects is largely empirical and requires optimization of the primary MB-FUS parameters: ultrasound intensity and MB dose. Treatment effects are determined by monitoring of bubble cavitation (e.g., acoustic emissions dose (AED)) signal during treatment (21), of successful BBB opening by extravasation of the magnetic resonance imaging (MRI) contrast agent gadolinium, of tissue damage on post-treatment MRIs, and of reported neurological events (3). However, second-order effects of BBB opening, including neuroinflammatory activation, systemic immune response, and behavioral changes, are challenging to assess, may develop more slowly, and remain incompletely characterized (22–24).

Expansion of MB-FUS in the clinical sphere is not without caveats; despite generally favorable safety profiling, isolated cases have been reported where intense FUS treatments have resulted in coma with prolonged neurobehavioral adverse effects (25) as well as less serious adverse effects including headache, syncope and postural hypotension (1). These caveats, together with the broad aim of expanding treatment indications for MB-FUS, require thorough systematic evaluation of the safety of a broad range of high intensity treatment parameters. Due to safety concerns and limitations in human studies, such a comprehensive evaluation is only possible in the preclinical sphere. Importantly, a holistic safety evaluation must test acute and long-term effects of MB-FUS treatments and go beyond common readouts (e.g., histology and simple behaviors such as the Irwin test (24, 26)) and explore effects on complex behaviors which require the interplay between brain areas.

Here, we describe a critical step towards a comprehensive safety evaluation of MB-FUS: we tested acute and long-term effects of a wide range of prescribed AEDs, both comparable to and higher than clinically used doses. We probed the effects not only on observable tissue damage, quantifying the readouts of multiple MRI sequences, but also on complex learned behavior in rats. For the latter, we leveraged a well-established, custom motor skill task in which rats develop idiosyncratic, spatiotemporally precise, and complex *de novo* movement patterns over several months of training (27). We determined, in rats performing at expert levels, how different MB-FUS dosing regimens affected their complex learned behavior. We took advantage of the highly sensitive behavioral readouts provided by our task, including not only performance measures, but also precise movement kinematics. Machine learning-aided pose estimation (28) enabled us to track the detailed learned movement patterns before and after treatment and to determine if and how the learned behavior was affected.

In addition, insights into the motor circuitry underlying the execution of this task from our previous work (27, 29, 30) allowed us to test whether observed behavioral impairments were specific to the targeted anatomical location or due to nonspecific, non-local effects of MB-FUS treatment. We previously found that the motor cortex, while necessary for motor skill learning, becomes dispensable with training, and execution of the learned skill is driven by subcortical circuits, most prominently the basal ganglia and thalamus (27, 29, 30). Comparison of cortical- and subcortical-targeting of MB-FUS treatment therefore allowed us to test the specificity of its effects.

By combining multiple MRI sequences, histology, and our sensitive behavioral task, we established a novel preclinical platform to test the acute and long-term effects of a broad range of MB-FUS dosing regimens. While all doses we tested induced BBB opening, we found a marked dose-dependent increase of adverse effects. Importantly, low dosing regimens did not cause overt adverse effects in our various readouts. However, higher doses, at the limit of or beyond the currently clinically used range, caused significant disruptions of both the targeted brain tissue and the probed behavior. Furthermore, behavioral impairments were specific to treatments targeting the relevant (subcortical) neural circuitry. Our longitudinal experiments also showed that some adverse effects persisted throughout our observation period of several weeks. The time course of recovery, however, differed substantially across readouts – while general behavioral performance measures recovered after about a week, tissue disruptions persisted for at least 3 weeks. Notably, even the highest-intensity treatment regimen produced markedly variable effects across individuals, with a subset of animals showing recovery of behavioral performance measures, but long-lasting significant changes in the quality of their learned movement patterns. Taken together, while low dose MB-FUS treatments appear safe across our readouts, higher intensity regimens pose notable risks. Variable effects across individuals and distinct time courses of recovery across multiple measures strongly underscore the need for a holistic evaluation across timescales, that goes beyond common readouts, to validate new MB-FUS regimens for safe future clinical applications.

## RESULTS

### Subspot-level AED provides a more complete description of MB activity than common measures

To broadly explore potential adverse side effects of MB-FUS treatment on both the behavioral and histological level, we exposed rats to various dosing regimens, covering a wide range of the relevant parameter space for FUS and MB concentration (combinations labeled ‘Low’, ‘Medium’, ‘High’, and ‘Ultra-High’) as well as different targeting depths (superficial (S; cortex), deep (D; cortex and subcortical areas), and intermediate (I)) to test for circuit-specificity (***Table 1;* *Fig. 1B***).

**Table 1.**
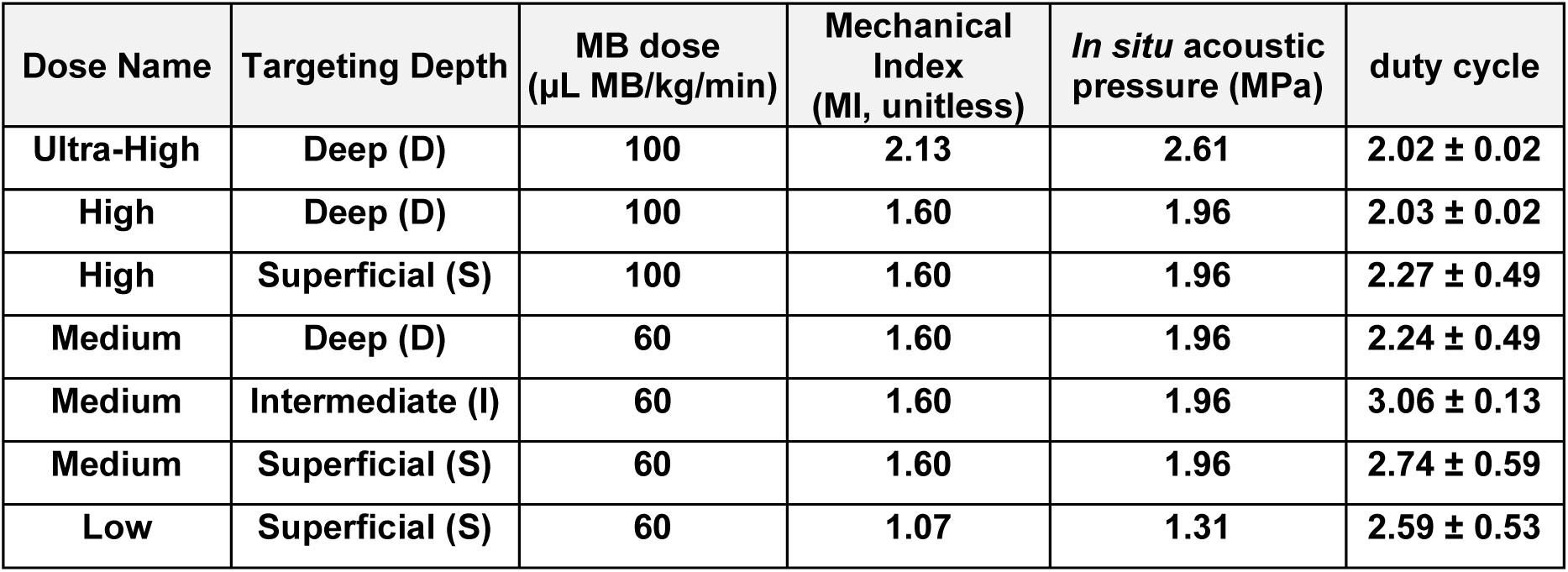
Applied treatments and FUS properties. Treatment doses were defined by device power and MB dose. Treatment depth was determined by transducer properties and confirmed with T1c MRI. Duty cycle is reported as average ± standard deviation of the most recent 3-5 treatments in each group.

| Dose Name | Targeting Depth | MB dose<br>(μL MB/kg/min) | Mechanical<br>Index<br>(MI, unitless) | *In situ* acoustic<br>pressure (MPa) | duty cycle |
| --- | --- | --- | --- | --- | --- |
| Ultra-High | Deep (D) | 100 | 2.13 | 2.61 | 2.02 ± 0.02 |
| High | Deep (D) | 100 | 1.60 | 1.96 | 2.03 ± 0.02 |
| High | Superficial (S) | 100 | 1.60 | 1.96 | 2.27 ± 0.49 |
| Medium | Deep (D) | 60 | 1.60 | 1.96 | 2.24 ± 0.49 |
| Medium | Intermediate (I) | 60 | 1.60 | 1.96 | 3.06 ± 0.13 |
| Medium | Superficial (S) | 60 | 1.60 | 1.96 | 2.74 ± 0.59 |
| Low | Superficial (S) | 60 | 1.07 | 1.31 | 2.59 ± 0.53 |

Importantly, to draw conclusions regarding the relationship between distinct treatment parameters and potential adverse effects, a reliable measure of the effectively applied dose is critical. Mechanical index (MI) values are a common measure of FUS dose and provide a useful description of prescribed acoustic exposure. However, recent studies have shown that local MB activity is influenced by more than just acoustic pressure and that the AED is a more informative predictor for the effects of MB-FUS treatment (2). Indeed, common measures were not able to fully distinguish our regimens: despite distinct MB concentrations and targeting, our Medium/S, Medium/I, Medium/D, High/S, and High/D conditions all had *in situ* acoustic pressures of 1.96 MPa and an *in situ* MI of 1.6. Only the Ultra-High/D condition showed higher values with an acoustic pressure of 2.61 MPa and a MI of 2.13 (***Table 1***). We note that this condition has an MI beyond the FDA’s limit for diagnostic ultrasound imaging at a MI of 1.9. While this is not a therapeutic safety threshold for MB-FUS BBB opening, we included the Ultra-High regimen to go beyond current clinical practice, to determine whether our novel combination of multimodal readouts could detect treatment-associated neurobehavioral, imaging, and histological changes, and to help define the boundary between therapeutic BBB opening and adverse biological effects.

To better distinguish the effects of our treatment regimens and to consider all relevant variables, we calculated the AED at the level of the individual subspots of the treatment grid for our four highest intensity regimens (***Figs. 1, 2***) (see Methods). This allowed us to determine the spatial distribution of acoustic effects over the treatment grid and rat brain. Indeed, we found that both the subharmonic AED (AED_SH_) and the ultraharmonic AED (AED_UH_) increased with higher estimated *in situ* pressure, higher MB dose, and deeper anatomical target location (***Fig. 2A,C***).

**Figure 2.**
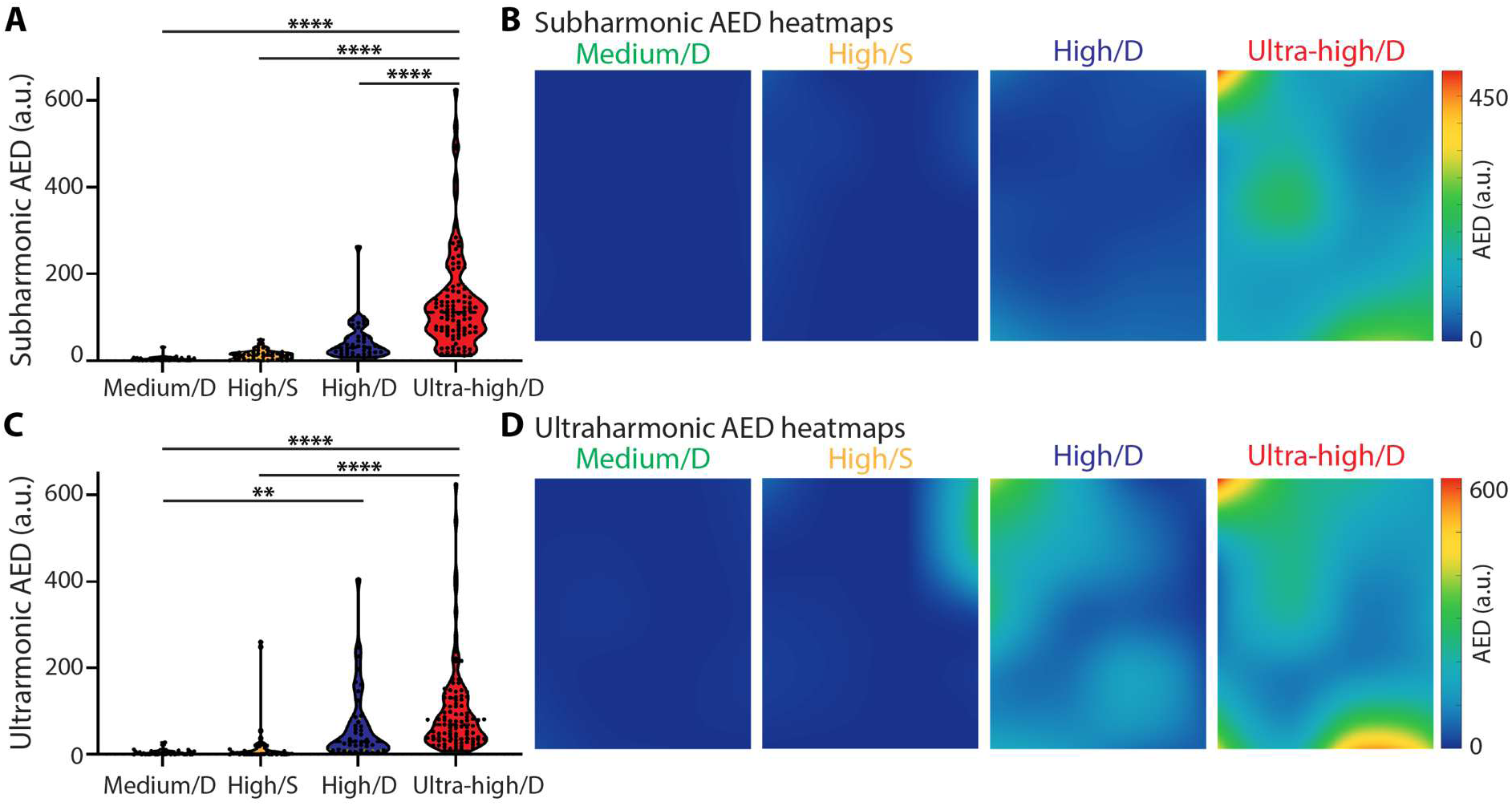
Subspot-level Acoustic Emissions Dose (AED) reveals acoustic pressure-,microbubble dose-, and target-dependent microbubble activity during MB-FUS. **A** Subspot-level AEDSH distributions. Dots represent treatment subspots. Subspot-level AED was calculated from acoustic emission recordings acquired during MB-FUS treatments. The following number of sonications were analyzed: 500 for Medium/D, 1,000 each for High/S, and High/D, and 2,500 for Ultra-High/D. **B** Representative interpolated AEDSH heatmaps, with AED values in arbitrary units. For corresponding median AEDs and interquartile ranges, see ***Supplementary Table S1. C,D*** As A,B for AEDUH. Statistics (A, C): 1-way ANOVA with Tukey’s multiple comparisons test. * p<0.05; ** p<0.01; *** p<0.001; ****p<0.0001.

The differential effects of the dosing regimens became even more apparent in spatial AED heatmaps, which showed heterogeneous MB activity within each treatment grid (***Fig. 2B,D***). The Medium/D dose showed low AED_SH_ and AED_UH_ across most subspots. Increasing the MB dose increased AED in a depth-dependent manner; the High/S regimen showed lower AED values than the High/D condition. The highest AED values and broadest spatial heterogeneity were observed in Ultra-High/D treatments (***Fig. 2B,D; Supplementary Table S1***). Together, this indicates that subspot-level AED captures local differences in MB activity that are not fully described by estimated pressure or MB dose alone. Target location also contributed to the detected acoustic emission response, supporting the use of subspot-level AED as a spatially resolved readout that can be compared directly with MRI-defined BBB opening and other downstream biological outcomes.

### MB-FUS treatment causes dose- and targeting-dependent acute disruptions of learned motor behavior

To determine the effects of our MB-FUS treatment regimens on the generation of complex learned behavior, we took advantage of our custom motor skill paradigm (see above; ***Fig. 3A***). In this task, rats gain water rewards for pressing a lever twice with a specific inter-press interval (IPI; target: 700 ms; ***Figs. 3A,B***) (27, 29, 30). Over weeks of operant shaping in individual, fully-automated training cages, rats develop *de novo* idiosyncratic movement patterns, including motor elements in-between the lever presses, allowing them to approximate the target IPI with increasing precision. The emerging movement patterns become increasingly complex, spatiotemporally precise, and stereotyped. Rats also learn to keep a required time-out period after unrewarded trials (inter-trial interval, ITI, >1.2 s) as part of the overall task structure. For a detailed characterization of the task and the development of the learned movement patterns, see (27).

**Figure 3.**
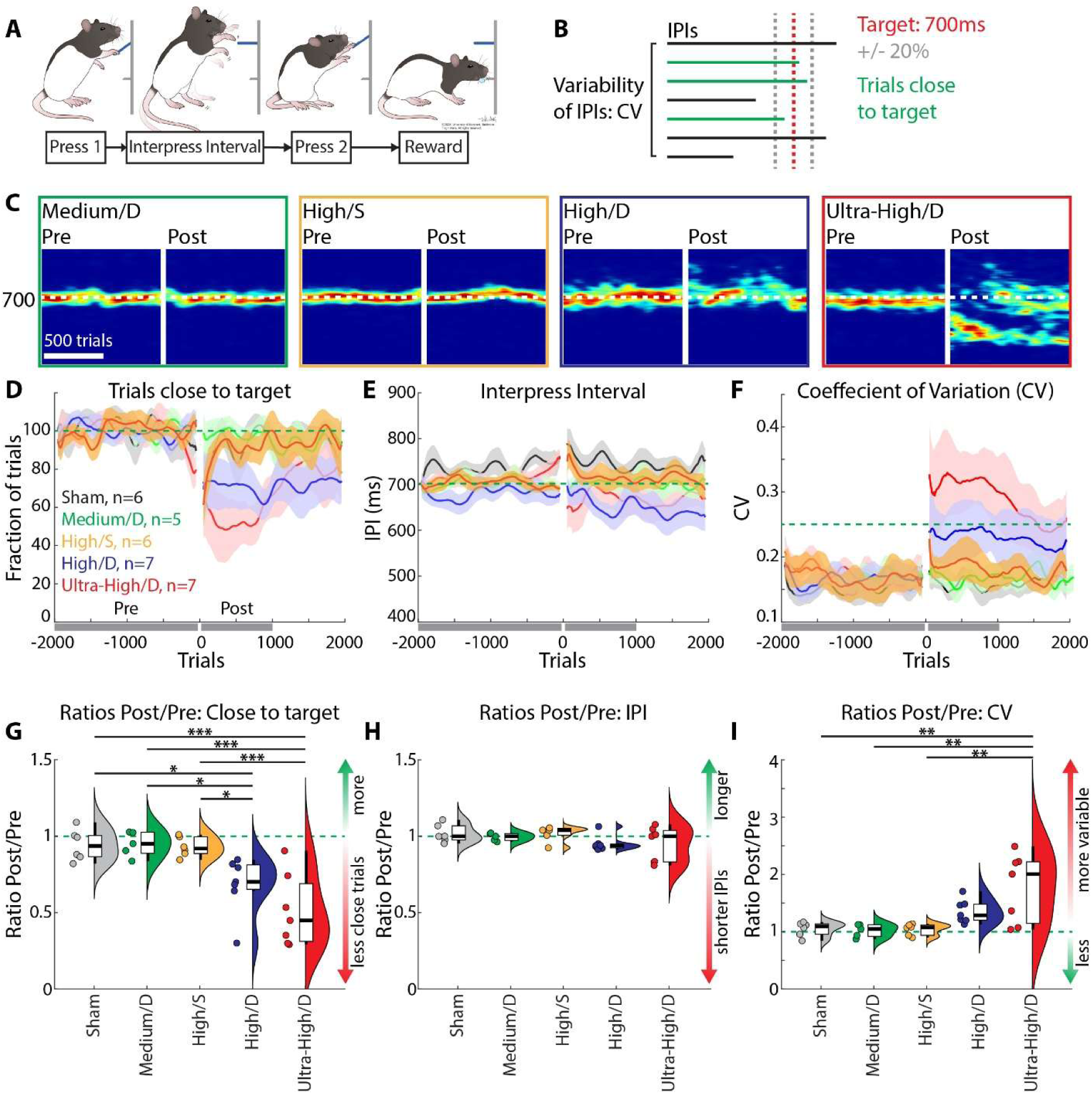
Acute effects of MB-FUS treatment on performance of a learned motor skill. **A** Motor skill paradigm. Rats learn to press a lever twice with an inter-press interval (IPI) close to a fixed target (700 ms) for water rewards scaled by distance to the target. Over thousands of trials, rats develop idiosyncratic movement patterns spanning the IPI. For details see text and (27). **B** Performance measures. We determine the fraction of trials with IPIs close to the target, i.e., within +/-20%, and IPI variability (Coefficient of Variation: CV). **C** Performance of example MB-FUS treated rats. Probability density plots show the likelihood of the generation of specific IPIs (1,000 trials pre- and post MB-FUS treatment. Warmer colors indicate higher likelihoods. **D** Population plot for the fraction of trials close to target pre- and post-MB-FUS and sham treatments, normalized to pre-treatment performance. Population means and SEM (for D-F). **E** Population plot for IPIs pre- and post-MB-FUS. **F** Population plot of the CV as measure of behavioral variability pre- and post-MB-FUS. **G** Ratios of the fraction of trials close to target between post- (1,000 trials) and pre- (2,000 trials) MB-FUS treatment (cf. D). Violin plots (Kernel Density) with boxplots (Whiskers: Minimum/Maximum values; box: 25^th^ and75^th^ percentile; line: Median) and dots as individual rats. **H** As G, but for IPI ratios. **I** As G, but for CV ratios. Statistics (G-I): 1-way ANOVAs with Post-hoc tests (Holm), see ***Supplementary Table S2*** for detailed results. * p<0.05; ** p<0.01; *** p<0.001.

We treated rats that performed at expert levels in this task (see Methods for expert criteria) with our broad range of MB-FUS regimens (***Table 1***). After recovery from anesthesia, rats were immediately returned to their training cages, allowing them to resume training the following night and to continue training for at least 15 days, providing insights into both acute and long-term effects. A detailed analysis of the performance of treated animals revealed dose-dependent acute disruptions of the learned motor behavior (***Fig. 3, Supplementary Fig. S1***). Rats treated with the highest dose (Ultra-High/D) showed significant disruptions in the first 1,000 trials after treatment when compared to sham-treated rats: their generated IPIs were more variable than during the pre-treatment baseline (Coefficient of Variation (CV) of IPIs; ***Fig. 3F, I***), leading to a severe reduction in the fraction of trials close to the target IPI (trials in the range of 700 ms +/− 15%; ***Fig. 3D, G***), and an alteration of their IPI distributions (***Supplementary Fig. S1B***). Furthermore, these rats showed disruptions in their ability to adhere to the overall task structure: they generated less trials with exactly two lever presses (***Supplementary Fig. S1D-F***) and their learned, distinct IPI and ITI distributions (around the target 700 ms interval vs. timeout after unrewarded trials >1.2 s), became more similar (***Supplementary Fig. S1C***).

Notably, while treatment with the second-highest dose (High/D) also caused a significant, albeit less pronounced, acute decrease in performance (***Fig. 3, Supplementary Fig. S1***), lower treatment doses (Medium, ***Fig. 3, Supplementary Figs. S1, S2***; Low, ***Supplementary Fig. S2***) did not cause acute performance deficits. This suggests that MB-FUS treatment can acutely disrupt complex learned motor behavior in a dose-dependent manner.

A critical question regarding potential adverse effects of MB-FUS treatment is whether they are due to localized disruptions of the targeted neuronal circuitry or due to nonspecific, potentially brain-wide effects. To approach this, we took advantage of our previous findings on the neural circuitry involved in our task: Skill execution in experts requires subcortical areas, prominently including the basal ganglia and thalamus, but is independent of (motor) cortex (27, 29, 30). We therefore adjusted our MB-FUS treatments to preferentially affect either the cortex or both the cortex and subcortical areas by varying the targeting depth (superficial (S), intermediate (I), deep (D); ***Table 1***; see Methods). In line with our previous findings, we observed a stark contrast between superficial and deep targeting in our ‘High’ dosing regimen: While deep targeting (High/D) significantly disrupted task performance (see above), superficial targeting (High/S) had no acute effect on performance (***Fig. 3, Supplementary Fig. S1***), supporting target-dependency instead of non-localized effects. Notably, at lower doses, no effects on performance were observed, independent of targeting depth (***Supplementary Fig. S2***). Together, our findings suggest that adverse effects of MB-FUS are both dose- and target-dependent.

### MB-FUS treatment causes dose- and targeting-dependent acute BBB opening and histological changes

To broadly visualize the effects of our MB-FUS treatments on the brain and gather a range of clinically relevant information, we performed histological analyses, analyzed multiple MRI sequences and, beyond common practice, quantified the observable effects of MB-FUS. T2w and T1w sequences provide structural anatomy and high-resolution morphology. T2w images show water content in tissue; areas of edema or inflammation will be bright. T1w images, in which fluid appears dark, were used to assess anatomy and as negative control for contrast-enhanced images. Our main metric for successful BBB opening was enhancement of the T1-contrast (T1c) sequence, i.e., the T1w sequence after injection of the MRI contrast agent gadolinium, which shows areas with BBB opening as bright regions. SWI sequences are highly sensitive for molecules that distort the local magnetic field, such as iron and blood, allowing us to quantify MB-FUS-induced microhemorrhage (20). While this sequence is not commonly used preclinically for BBB opening studies, we included it to explore acute and long-term structural damage.

We focused our analysis on the four highest-intensity regimens (Ultra-High/D, High/D, High/S, Medium/D), which caused dose- and targeting-dependent behavioral effects (***Fig. 3***). T1c images taken directly after MB-FUS treatment in both trained and naïve animals and quantification of T1c enhancement revealed BBB opening in all imaged treatment regimens, largely restricted to the targeted region (***Fig. 4A,B; Supplementary Fig. S3***). Notably, we found differences in the location of the BBB opening: while deep targeting led to widespread opening throughout the hemisphere, including both cortical and subcortical areas (Medium/D, High/D, Ultra-High/D; ***Supplementary Fig. S4B***), superficial targeting caused BBB opening only in cortical areas (High/S; ***Supplementary Fig. S4B***). This was in line with the lack of behavioral disruptions in the High/S group (***Fig. 3***) and the need for subcortical, not cortical circuits for task execution (27, 29). Additionally, we assessed the presence of microhemorrhages and edema using SWI and T2w sequences, respectively. SWI imaging and quantification revealed little magnetic field disruption in Medium/D and High/S treatments, but increased levels in the High/D and Ultra-High/D groups compared to sham-treated animals (***Fig. 4C***). Upon visual inspection, T2w scans showed little to no edema across all doses on the day of treatment (***Fig. 4A***).

**Figure 4.**
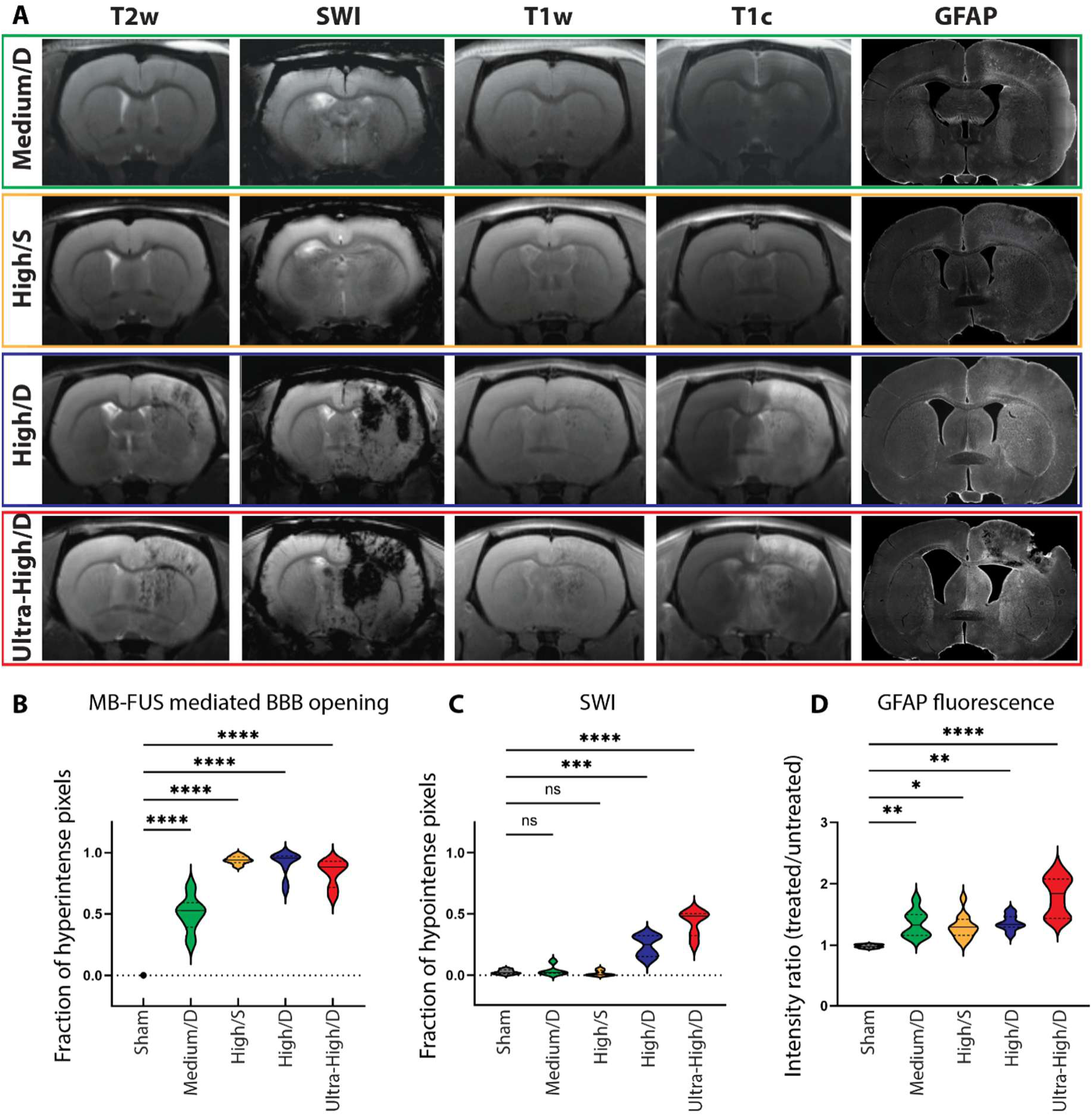
Imaging and histological readouts for high-intensity MB-FUS dosing regimens. **A** MRI sequences and GFAP fluorescent staining from example rats, captured within 30 minutes or 72 hours, respectively, after MB-FUS treatments. **B** Population quantification of T1c MRI sequence as a marker of BBB opening: Proportion of contrast-enhancing pixels within the respective treatment ROI across imaging planes. **C** Population quantification of SWI MRI sequence as an indicator for potential microhemorrhages: Proportion of hypointense pixels within the respective treatment ROI across imaging planes. **D** Population quantification of GFAP fluorescent staining as a marker for astrocyte activation: Ratio of fluorescence intensity between treated and untreated hemispheres across six planes. Statistics: 1-way ANOVA with multiple comparisons corrections. Only comparisons to the sham group are shown, for other comparisons, see ***Supplementary Table S5***. * p<0.05; ** p<0.01; *** p<0.001; **** p<0.0001.

In addition to these MRI sequences, we characterized MB-FUS treatment effects histologically. We focused on GFAP (Glial Fibrillary Acidic Protein), a marker for astrocyte activation in response to damage and/or inflammation in the CNS (31). We treated naïve, untrained rats with MB-FUS and after three days, allowing for an astrocyte response (32), fixed and extracted the brains. Subsequently, immunofluorescent staining revealed significantly increased GFAP levels compared to sham-treated control rats across all conditions (measured as ratio of GFAP levels in treated vs. untreated hemispheres) (***Fig. 4A,D***), indicating prolonged astrocyte activation after MB-FUS treatment. Notably, GFAP levels were also dose-dependent, with higher levels in the Ultra-High/D group compared to the other tested groups (***Fig. 4A,D; Supplementary Fig. S4D***).

### MB-FUS treatment causes dose-dependent long-term disruptions of learned behavior

To determine whether the acute behavioral deficits in our Ultra-High/D and High/D groups (***Fig. 3***), persisted over extended periods of time, we continued to train treated animals for at least 15 days. To analyze their behavior over weeks, we averaged the performance across the animals’ two training sessions every night to provide daily measures. Compared to sham-treated animals, rats in the High/D group showed significant behavioral impairments on Day 1 after treatment with a reduced fraction of trials close to the target (***Fig. 5A,B***), due to the generation of both shorter and more variable IPIs (***Supplementary Fig. S5***). However, the High/D animals quickly recovered, and on Day 2 their performance was not significantly different from sham animals (***Fig. 5A,B; Supplementary Fig. S5***). In stark contrast, rats in the Ultra-High/D group showed prolonged behavioral deficits: After a similarly strong reduction in the fraction of trials close to the target on Day 1 (***Fig. 5A,B***), they only slowly recovered, showing significant differences compared to sham-treated controls until Day 9 after treatment (***Fig. 5A,B***). These deficits were partially driven by a severe reduction in task engagement. On Day 1, most rats did not participate in the task at all and continued to miss more than 20% of training sessions until Day 5 (***Fig. 5C,D***). In contrast, after High/D treatment, animals only missed sessions on Day 1 (***Fig. 5C,D***). Notably, animals in the Ultra-High/D group which did perform the task on the first day also showed less behavioral impairment overall compared to animals that did not perform initially (***Fig. 5, Supplementary Fig. S5***). Taken together, our results suggest that MB-FUS treatment can cause long-term deficits in learned motor behavior for more than 1 week after treatment.

**Figure 5.**
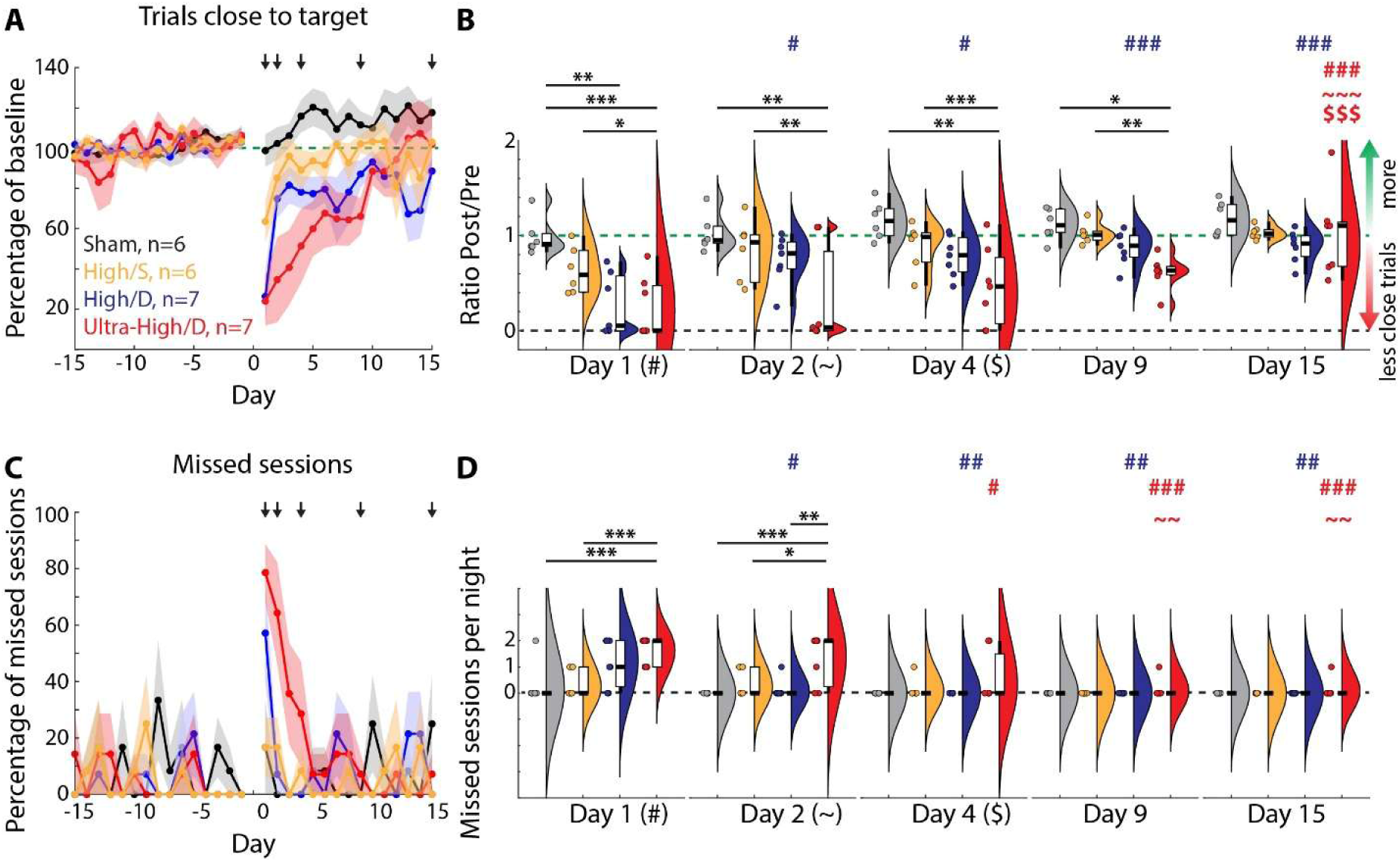
Long-term effects of MB-FUS treatment on motor skill performance. **A** Trials close to target across days pre- and post-MB-FUS treatment, normalized to pre-treatment performance. Daily performance is averaged across two daily sessions. Missed sessions are considered as having a fraction of trials close to target of 0. Dots: Population average per day. Error bars: SEM. **B** Ratios of the fraction of trials close to target between post- and pre-treatment (averaged across 10 days) for selected days. Violin plots with boxplots and individual rats as in Fig. 3G. C Like A, but for fraction of missed sessions per day. Rats can perform in up to two sessions per night. Sessions are counted as missed if no trials are performed during session time. **D** Like B, but for number of missed sessions per night. Statistics (B,D): Linear Mixed Model with planned contrasts (Comparison across groups for each day and comparisons across days for each group). For detailed results see ***Supplementary Table S7*.** Stars indicate significant differences across groups on the same day. Symbols indicate significant differences across days for a given group (Symbol indicates compared day (e.g., # compared to Day 1), color indicates group (e.g., red for Ultra-High/D). * p<0.05; ** p<0.01; *** p<0.001.

To determine whether treatments with lower MB-FUS doses may cause delayed or long-term behavioral deficits, we also analyzed the performance of the low-dose groups throughout the same 15-day period (***Supplementary Fig. S6***). Animals in the Medium/S and Medium/I groups showed some effects on performance on Day 1 (reduced fraction of trials close to the target; ***Supplementary Fig. S6A,B***), partially due to an increase in the number of missed sessions (***Supplementary Fig. S6A,B***). However, these initial deficits were resolved by Day 2, and no additional deficits emerged over the 15-day post-treatment period.

### MB-FUS can disrupt the detailed kinematics of learned motor skills

Our motor skill paradigm provides detailed readouts of the fine-scaled, highly stereotyped kinematics our animals develop over weeks of training, allowing us to probe the effects of MB-FUS on learned behavior at a level of detail beyond typical performance measures. Notably, potential kinematic changes may indicate impairments of precise motor execution and/or of the previously formed motor memories. Machine learning-aided pose estimation (28) allowed us to track the animals’ hand movements during and in-between the lever presses in the task, before and after MB-FUS treatment (***Fig. 6***). To determine the effects of our treatments on the stereotypy and structure of the complex learned movement patterns, we calculated the trial-by-trial correlations across movement trajectories separately before treatment, on the 1^st^ active day after treatment (i.e., the 1^st^ day with at least 20 trials; ‘Day 1’) and on Day 15 after treatment. On Day 1, both the High/D and the Ultra-High/D groups showed significantly lower trial-by-trial correlations than before treatment (***Fig. 6A,B***), and lower correlations than sham-treated animals on Day 1 (***Fig. 6B***). This suggests that high intensity MB-FUS treatments caused an impairment of the learned stereotypy and increased movement variability. However, these effects were transient, and the trial-by-trial correlations returned to pre-treatment levels on Day 15 (***Fig. 6B***). In line with our results on task performance, low intensity MB-FUS treatments had no effect on the stereotypy of the learned kinematics, with the exception of transiently decreased correlations after the Medium/S treatment (***Supplementary Fig. S7***).

**Figure 6.**
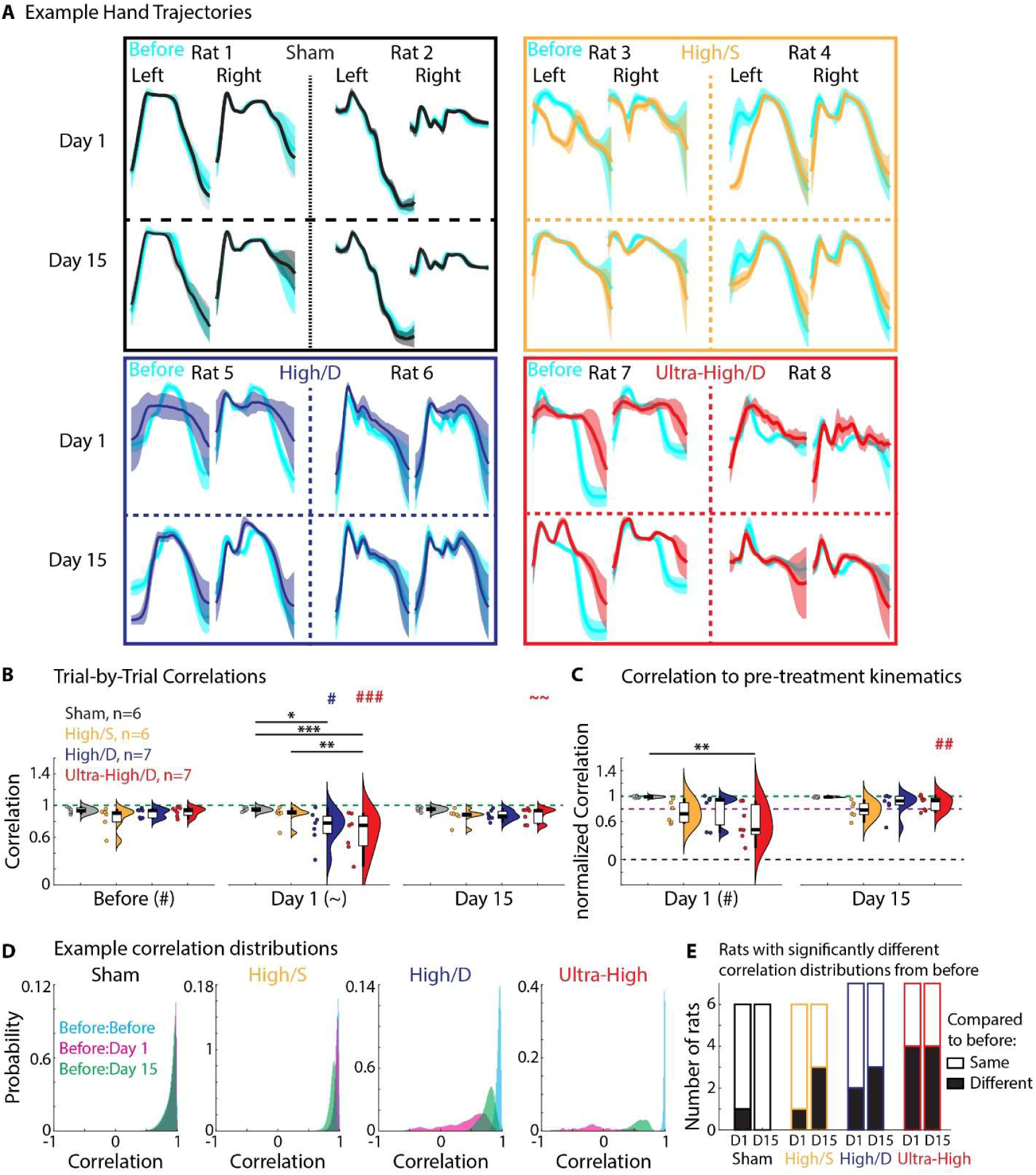
Acute and long-term effects of high-intensity MB-FUS treatment on complex learned motor skill kinematics. **A** Hand movement trajectories (vertical dimension; left and right hands) of example rats before and after MB-FUS treatments. Cyan and colored traces show pre- and post- treatment trajectories, respectively. Shown are mean trajectories +/− SEM across all trials with the dominant movement mode (see Methods) on a given day. **B** Trial-by-trial correlations across all trials before MB-FUS treatment (across 5 days), on the first day of performance after treatment (Day 1) and on Day 15 after treatment. **C** Correlations between trajectories pre- and post-MB-FUS treatment, normalized to the mean trial-by-trial correlation pre-treatment for each animal (see B). For B, C: Violin plots with boxplots and individual rats as in Fig. 3G. Statistics: Linear Mixed Model with planned contrasts (Comparison across groups for each day and comparisons across days for each group). For detailed results see ***Supplementary Table S10***. Stars and symbols indicate significance as in Fig. 5. **D** Trial-by-trial correlation distributions for multiple comparisons (Before to Before, Before to Day 1, Before to Day 15) for example rats. Note different y-axis scales. **E** Number of rats with and without correlation distributions on Day 1 and Day 15 that are significantly different from Before MB-FUS treatment (see D), as determined by KS-tests for each individual rat. * p<0.05; ** p<0.01; *** p<0.001.

The transient loss of stereotypy after high-intensity MB-FUS treatment might be due to motor deficits, preventing rats from generating precise kinematics, or due to disruptions of formed memories (or their recall), impairing the generation of what had previously been learned and requiring animals to find new, distinct movement solutions. To distinguish these possibilities, we directly compared the movements before and after treatment (***Fig. 6C-E***). First, we calculated the correlations between movement trajectories before and after treatment (normalized by the average correlation of the pre-treatment trajectories; ***Fig. 6C***). When we compared the treatment groups, we found that the Ultra-High/D group showed significantly lower before-to-after correlations than sham-treated rats on Day 1 (***Fig. 6C***), indicating that these rats generated movement patterns distinct from the previously learned ones. On average, however, the before-to-after correlations on Day 15 were not different anymore between Ultra-High/D and sham-treated animals (***Fig. 6C***).

Importantly, however, while the group results suggest transient effects on the learned kinematics, more consistent with transient motor deficits, the movement patterns of a subset of animals across groups remained notably distinct from their initially learned movements, even on Day 15 (***Fig. 6A***). These rats showed markedly low before-to-after correlations throughout our observation period with values below 0.8, while all sham-treated animals had before-to-after correlations of at least 0.95 (***Fig. 6C***; see dashed line at 0.8), suggesting that the treatment effects were variable and that some animals suffered from prolonged behavioral changes. To further investigate this, we directly compared the distributions of the before-to-before, before-to-Day 1 and before-to-Day 15 correlations for individual animals (***Fig. 6D,E***). If the movement patterns of an animal were changed after MB-FUS treatment, these distributions should be significantly different. Indeed, as the example animals in ***Fig. 6D*** show, while sham-treated animals had highly similar distributions, individual animals across the treatment groups had significantly different distributions. Notably, the higher the treatment dose was, the more animals showed significantly different correlation distributions, with more than half on both days 1 and 15 in the Ultra-High/D group (***Fig. 6E***). Taken together, this strongly suggests that a subset of animals in each treatment group changed their previously learned kinematics and adapted new, distinct movement patterns after MB-FUS treatment. Importantly, these changes could not be explained merely by the increased movement variability (***Supplementary Fig. S8A***): We tested whether the variability of the movements of individual animals after treatment (trial-by-trial correlations on Day 1 or Day 15; cf. ***Fig. 6B***) was correlated with the similarity of the before and after treatment movements (before-to-after correlations; cf. ***Fig. 6C***). These measures were mostly uncorrelated, and even for the Ultra-High/D group, we only detected a correlation on Day 1, not on Day 15 (***Supplementary Fig. S8A***). Furthermore, we also found that the before-to-after correlations did not correlate with the performance of the individual animals (***Supplementary Fig. S8B***; Post/Pre ratio of the trials close to the target; cf. ***Fig. 5A***). Taken together, these results indicate that a subset of animals was able to regain performance in the task and to generate stereotyped movements after MB-FUS treatment, but that these movement patterns had significantly changed compared to the ones they had initially learned.

### High intensity MB-FUS treatment causes long-term brain tissue disruptions

Since treatment with the highest intensity MB-FUS regimen (Ultra-High/D) caused prolonged impairments across our behavioral performance measures for about one week (see above; ***Fig. 5***) and affected the kinematics in some animals for even longer (***Fig. 6***), we decided to determine whether the time course of recovery from the acute brain tissue disruptions we observed (see above; ***Fig. 4***), matched the time course of recovery for behavioral deficits. For this, we collected a series of MRIs over the course of 21 days after Ultra-High/D treatment, focusing on the resolution of the observed acute microhemorrhages (***Fig. 4***) and, additionally, on the potential development and resolution of edema (33). SWI images indicated that magnetic field distortion (suggesting microhemorrhages) was high on the day of treatment, decreased significantly after 24 hours, and stabilized at a slightly increased level approximately 7 days after treatment (***Fig. 7A,C***), suggesting that the initial acute microhemorrhages partially resolved, but did not fully disappear over the course of 3 weeks, outlasting the observed behavioral performance deficits (***Fig. 5***). Similarly, T2w imaging suggested development and extended presence of fluid in the brain with a distinct time course. While edema was almost undetectable on the day of treatment, it developed over the course of 24 hours and remained, with some fluctuations, at comparably high levels over the course of the probed 21 days (***Fig. 7B,D***). The morphology of physiological effects and presence of fluid pockets in the brain may be suggestive of gliosis and prolonged neuroinflammation (34).

**Figure 7.**
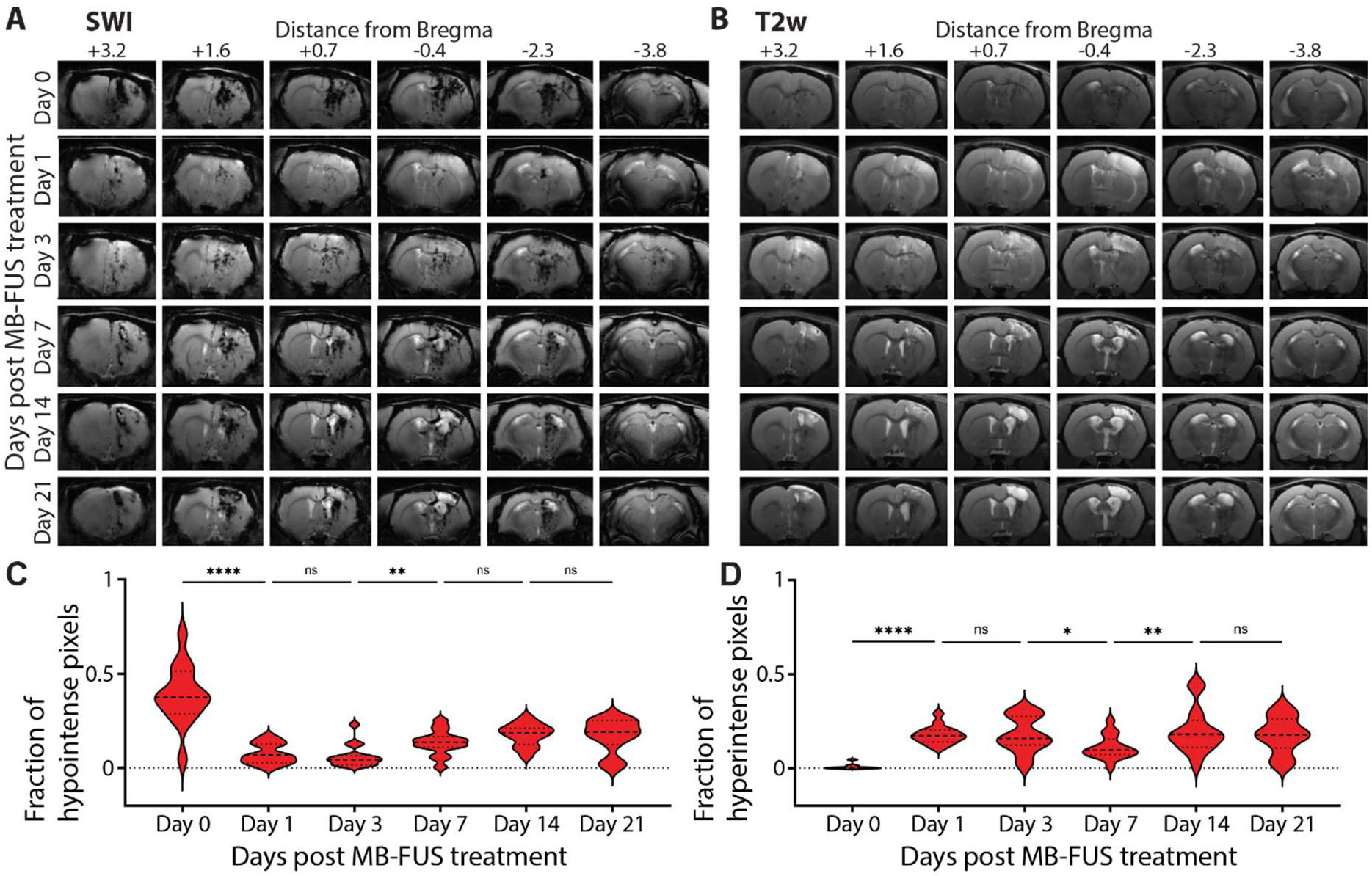
Longitudinal imaging after MB-FUS treatment with the Ultra-High/D dose. **A** SWI and **B** T2w images of an example rat over 3 weeks after Ultra-High/D MB-FUS treatment across multiple imaging planes. C, D Population quantifications for longitudinal SWI and T2w sequences, respectively. Values are averaged for each rat (n=3) across 5 imaging planes. Statistics: 1-way ANOVA with multiple comparisons corrections. Shown are only comparisons between consecutive days; additional comparisons shown in ***Supplementary Table S12.*** * p<0.05; ** p<0.01; *** p<0.001; **** p<0.0001.

## DISCUSSION

While MB-FUS is a highly promising and versatile emerging treatment approach across a range of central nervous system disorders, defining conditions that maximize BBB opening without inducing adverse biological effects remains a fundamental challenge. Addressing this challenge is becoming increasingly urgent due to rapidly growing clinical and preclinical interest in expanding MB-FUS use cases. In this study, we take a significant step forward by establishing a novel, multimodal platform to test the effects of MB-FUS on brain structure and function as well as on behavior across dosing regimens; we combine MRI, histological assessment, and a highly sensitive assay of complex learned behavior in rats. Importantly, while our results confirm that low intensity MB-FUS treatment appears safe across our readouts, we show that MB-FUS can cause acute (***Figs. 3, 4***) and long-lasting (***Figs. 5-7***) adverse effects that are strongly dependent on both the prescribed dose and targeting of the treatment. Although BBB opening was achieved in all regimens (***Fig. 4, Supplementary Fig. S3***), only the higher-intensity deep-targeting FUS settings caused behavioral impairments (***Figs. 3, 5***), alterations of movement kinematics (***Fig. 6***), and pronounced tissue injury (***Fig. 4***). Furthermore, our multimodal readouts revealed different time courses of the development and recovery of adverse effects (***Figs. 5-7***) as well as marked inter-individual variability (***Figs. 5, 6***), suggesting complex relationships between behavioral and brain tissue disruptions. Together, these results do not only suggest that expansion of MB-FUS to higher dosing regimens demands caution, but also that common acute, radiological readouts are insufficient to define a comprehensive safety profile, underlining the need for a multimodal assessment platform.

These results also complement recent clinical MB-FUS studies and emphasize the need for caution when comparing preclinical fixed-output exposures with clinical acoustic emission-controlled treatments. A Phase 1/2 trial of FUS in glioblastoma patients found that MRI-guided transcranial MB-FUS with adjuvant temozolomide enhanced drug delivery, with safety and feasibility as primary outcomes. BBB opening was observed in all treatments, and no treatment-related deaths were observed (1, 2). At the same time, recent experience in the broader FUS field underscores that even low-intensity FUS applications require rigorous characterization of exposure, monitoring, and longitudinal safety assessment. A single-patient adverse event report described brain injury during FUS-mediated neuromodulation for substance use disorder (25). That report is not directly comparable to the present MB-FUS study as it involved a different indication, target, device context, and treatment paradigm, but highlights nonetheless the translational importance of sensitive functional and imaging readouts when FUS exposures approach or exceed conventional safety reference points.

### Analysis of acoustic emissions provides a clinically comparable readout of dose-dependent effects of MB-FUS treatments

While fixed acoustic exposure parameters, like the estimated *in situ* MI, are important descriptors of MB-FUS treatments, AED analysis provides a more complete and biologically relevant treatment-dose metric, directly assessing the MB activity that produces the biological effect. Indeed, our AED results indicate that MB activity was influenced by more than acoustic pressure alone. At the same estimated *in situ* pressure and MB dose, cortex-targeted ‘High/S’ treatments produced lower AED_SH_ and AED_UH_ than striatum-targeted ‘High/D’ treatments. We also found that increasing the estimated *in situ* pressure in the Ultra-High/D regimen not only amplified MB activity, but also the spatial heterogeneity within the treatment grid (see the wide interquartile range (IQR); ***Supplementary Table S1***). Together, these findings emphasize that the effective acoustic emission response and biological effects of MB-FUS depend not only on the prescribed acoustic pressure, but on the local interaction between pressure, MB dose and dynamics, acoustic propagation, vascular density, tissue context and target location as well as on the detector’s sensitivity to acoustic emissions from each region. Subspot-level AED therefore provides a more direct readout of the local acoustic activity that drives BBB opening and possible biological effects.

We summarized our data at the subspot level not only to best capture these spatial effects, but also to allow comparison at the smallest unit most closely aligned with treatment delivery between preclinical and clinical treatments. In the related clinical acoustic emissions analysis of MB-FUS BBB opening, target-level AED was calculated from subharmonic emissions within each target, yielding an overall median AED of 0.58 a.u. with an IQR of 0.27-1.21 a.u. (2). The present preclinical values should not be interpreted as equivalent to clinical AED because the systems differ in frequency, detector geometry, acoustic path, skull transmission, and calibration. However, the workflow is conceptually aligned because it quantifies local MB activity at the level of individual treatment units and then allows those values to be compared with MRI-defined BBB opening.

In addition to the subharmonic AED emphasized in the clinic, we also quantified ultraharmonic AED. This additional readout is critical because ultraharmonics reflect nonlinear MB oscillation behavior and may provide complementary information about settings approaching stronger cavitation. Recent data-driven MB-FUS work further supports this concept by showing that acoustic emission features, including ultraharmonic components, can help identify patterns associated with the transition from stable to unsafe MB activity, thereby widening the treatment window for therapeutic delivery and biomarker release (i.e., tumor derived DNA) while limiting broadband events (35). AED_SH_ remains most directly aligned with the clinical dosing paradigm, while AED_UH_ provides additional preclinical information on nonlinear MB behavior, spatial heterogeneity, and the transition to higher-risk exposure settings. Together, the use of estimated *in situ* pressure, subspot-level AED_SH_, AED_UH_, and broadband safety monitoring provides a more complete description of MB-FUS exposure than any single metric alone.

### Multimodal, longitudinal readouts are necessary for a comprehensive MB-FUS safety profile

Our multimodal longitudinal readouts revealed that adverse effects can manifest differentially at various levels. Integration of behavioral and imaging measures painted a comprehensive picture of MB-FUS effects, which may be missed by common individual readouts. Our behavioral task combines a range of performance measures with detailed readouts of the generated complex movement patterns. This allowed us to dissociate a recovery of general performance deficits from persistent changes in fine-scaled learned behavior (***Figs. 5-6***). Similarly, our combination of multiple MRI sequences and, going beyond common practice, the quantitative comparison of MB-FUS effects over time (***Fig. 7***), revealed differential effects and clinically relevant insights. While the T1w contrast MRI sequence confirmed acute BBB opening across treatments (***Figs. 4, S3***), the SWI and T2w sequences revealed distinct time courses of microhemorrhage and edema development and recovery, outlasting BBB opening and behavioral effects (***Fig. 7****; see below*). These differences confirmed that relying only on individual measures would have produced incomplete safety profiles for our MB-FUS treatments.

### Adverse effects are dose- and targeting-dependent

Our combination of readouts revealed that adverse effects of MB-FUS treatment were specific and not merely consequences of anesthesia, handling, or generalized nonspecific BBB opening. We not only show a strong dose-dependency (***Figs. 3-6***), but also, by taking advantage of our insights into the task-related neural circuitry (27, 29, 30), a striking target-dependency: the same MB-FUS dose caused behavioral impairments when cortical and subcortical circuits, but not just cortex by itself, were targeted (***Figs. 3-6***). Taken together, these observations link behavioral impairments to severe disruptions of task-relevant circuitry rather than to diffuse brain-wide effects of MB-FUS. They also emphasize that treatment location is not a secondary technical detail. It is a critical part of the biological dose delivered to the brain.

### Acute adverse effects differ across readouts

Taking advantage of our multiple readouts allowed us to dive deeper into the differences between acute effects. First, while all treatments induced BBB opening (***Fig. 4, Supplementary Fig. S4***), only higher intensity groups suffered from acute tissue damage and behavioral deficits (***Figs. 3-4***), suggesting that BBB opening per se is not a reliable predictor for adverse effects. Similarly, the level of astrocyte activation was not indicative of other effects: despite comparable levels in the Medium/D, High/S and High/D groups (***Fig. 4A,D***), only the latter had notable acute microhemorrhages (***Fig. 4A,C***) and behavioral deficits (***Fig. 3***). Even our behavioral measures captured distinct effects: e.g., neither reductions in movement stereotypy (***Supplementary Fig. S8*A**), nor in behavioral performance (***Supplementary Fig. S8B***) of individuals on Day 1 were correlated with how much their movements changed after treatment.

### Longitudinal measures reveal increasing divergence of effects across readouts

The need to consider multiple readouts to define a holistic safety profile was further underscored by the longitudinal effects of MB-FUS treatments. First, similar acute effects could diverge over time: For example, despite comparable acute behavioral performance deficits in our two highest treatment groups (***Fig. 3***), the deficits only persisted in the Ultra-High/D group (***Fig. 5***). Second, we revealed striking dissociations of the development of the adverse effects across our readouts. For example, while behavioral performance deficits in the Ultra-High/D group were largely resolved after about 1 week (***Fig. 5***), notable changes in the learned movement patterns persisted in a subset of animals for at least two weeks (***Fig. 6***), and evidence of both microhemorrhages and edema persisted for at least 3 weeks (***Fig. 7***), strongly suggesting that tissue and functional recovery may proceed on, at least partially, independent time scales. These results have additional implications. While the emergence of edema is consistent with previous reports of increased markers of sterile inflammation at the proteomic and transcriptomic levels 24h after MB-FUS treatment [35], our longitudinal imaging strongly suggests that these changes can persist over extended periods. Another implication of the differential recovery of behavioral and tissue disruptions is that either the directly affected subcortical circuits or the circuits in the non-affected hemisphere have the capacity to eventually compensate for persisting tissue changes and to generate the previously learned, or at least stereotyped (***Fig. 6***), successful behavior (***Fig. 5***).

Importantly, the differing recovery time courses for tissue changes and behavioral impairments also highlight important practical points for safety assessment: First, no single measure is sufficient to fully describe the effects of MB-FUS treatments. Second, imaging at a single acute time point may underestimate transient but impactful impairments in circuit function, whereas persistent radiographic changes do not necessarily imply that overt behavioral impairment persists. A comprehensive evaluation of MB-FUS safety, therefore, requires both functional and structural readouts across extended time periods. The platform we present here is a first step towards establishing a holistic set of standardized readouts.

### Detailed dose characterization may explain intra-group variability of effects

The variability observed across animals and across outcome measures further supports the need for dose characterization with acoustic emissions. Marked inter-individual variability was evident in behavioral recovery, movement kinematics, and MRI findings (***Figs. 5-7***). This variability is consistent with the broader clinical experience in MB-FUS, where similar nominal treatment parameters can produce variable BBB opening across patients, targets, and treatment sessions. The ability to quantify subspot-level AED may help explain part of this variability by providing a direct readout of microbubble activity at the location where biological effects are expected to occur. In future studies, combining AED maps with longitudinal MRI and behavioral readouts could help distinguish animals with similar prescribed exposure but different effective microbubble activity, and could support more rational adjustment of treatment parameters across repeated sessions.

### Limitations

Despite our multimodal readouts and broad range of tested MB-FUS dosing regimens, our study is only a first step towards establishing standardized procedures to determine a comprehensive safety profile for MB-FUS across use cases and, possibly, for other emerging therapeutic approaches. Due to the limitations here, future studies should, for example, explore whether lower dose treatments that do not cause overt behavioral deficits may induce long-term brain tissue alterations. To directly test the relationships between long-term tissue changes and prolonged behavioral impairments, it will also be necessary to perform longitudinal imaging and behavioral testing in the same animals across dosing regimens. In addition, the mechanisms underlying the progressive tissue responses following our highest-dose MB-FUS treatment (***Fig. 7***) need to be further characterized to determine whether they arise from inflammatory, vascular, and repair-related processes and to understand how they may be mitigated. Our study also had technical limitations: While our motor learning paradigm provides high-dimensional, detailed behavioral readouts of a complex learned behavior, it does not address deficits in other behaviors or forms of learning, which may be differentially affected by MB-FUS treatment. Finally, due to hardware limitations of our single-element FUS transducer and the resulting focal dimension of the treatment field, our treatments lacked the anatomical precision to target individual brain structures. Importantly, however, the differential effects of superficial and deep targeting at the same dose are consistent with our previous insights into the underlying circuitry – targeting only of cortex, which is not necessary for task execution, had no effect, while targeting of cortex and the necessary subcortical structures disrupted behavior (***Figs. 3,5,6***). We note, however, that differential behavioral effects might also be due to a larger affected area in the deep, compared to the superficial treatment (*cf.* ***Fig. 4***), leading to an overall higher effective ‘dose’ and more severe effects. More localized targeting in future studies will allow us to address this caveat.

### Conclusions

Our findings are directly relevant to the broader translational trajectory of MB-FUS. Clinical studies have demonstrated that acoustic emissions-guided MB-FUS can achieve localized, controlled BBB opening and can be integrated with therapeutic strategies to improve drug delivery to the brain (1, 36, 37). Our results complement these clinical findings by defining functional safety boundaries in a preclinical setting with substantially greater experimental control. In this context, the present work supports a translational platform in which MB-FUS treatments are evaluated not only for their ability to open the BBB, but also for their effects on circuit function, behavior, and tissue integrity. Such an approach will become increasingly critical as the field moves toward larger treatment volumes, serial treatments, and more aggressive regimens designed to enhance the delivery of larger or less brain-penetrant therapeutics (e.g., antibodies, cell therapies).

More broadly, this study establishes a new paradigm for safety assessment that goes beyond standard histology and simple behavioral screening. By integrating multimodal imaging, circuit-informed targeting, complex learned behavior, and quantitative kinematic analysis, we show that subtle, yet consequential adverse effects can be detected longitudinally. Our data support the notion that preclinical evaluation of MB-FUS regimens should include not only confirmation of BBB opening and the absence of overt injury, but also sensitive measures of functional outcome. Together, these results define dose- and target-dependent constraints on MB-FUS-mediated BBB opening and support the use of behavioral readouts to identify safer operating regimens for brain-directed FUS.

## MATERIALS AND METHODS

### Animals

The care and experimental treatments of all animals were reviewed and approved by the University of Maryland Institutional Animal Care and Use Committee. For behavioral experiments, subjects were female Long Evans rats, 3–6 months old at the start of training (*n =* 46; Charles River). Because the behavioral effects of our treatments could not be specified in advance of the experiments, we chose sample sizes that would allow us to identify outliers and validate reproducibility. Animals for which no BBB opening was observed after MB-FUS treatment were assigned to the sham group. For histological assessment, female Long Evans rats were treated with MB-FUS at 2–3 months of age (n = 30). Rats with failed MB-FUS treatment or extensive tissue damage preventing reliable quantification were excluded from histological analysis. The investigators were not blinded to the group allocation during experiments and outcome assessment.

### Microbubble formulation

MBs were prepared by dissolving 1,2-dipalmitoyl-sn-glycero-3-phosphatidylcholine, 1,2-dipalmitoyl-sn-glycero-3-phosphatidylethanolamine-polyethylene glycol-5000, and 1,2-dipalmitoyl-sn-glycero-3-phosphatidic acid (Avanti Polar Lipids, Alabama) in glycerol, propylene glycol, and phosphate-buffered saline (PBS) at a molar ratio of 82:10:8. A 1.5 mL aliquot of this solution was transferred to a 2 mL serum vial (Duran Wheaton Kimble, Germany), degassed using a vacuum manifold, and then the vial headspace was filled with perfluoropropane gas (Fluoromed, Round Rock, TX). Immediately prior to MB-FUS treatments, MBs were activated by agitating the vial for 45 seconds using a Vialmix shaker (Lantheus Medical Imaging, North Billerica, MA), resulting in an average concentration of 2.2-5×10^9^ MBs/mL.

### Microbubble-enhanced focused ultrasound BBB opening

We performed MB-FUS treatments using a customized single-element FUS system equipped with a stereotactic-guided platform (Image Guided Therapy, Pessac, France). For treatment planning, we used landmark registration to the rat brain atlas to localize superficial (e.g., cortex) and deep (e.g., striatum) brain targets (38). The single-element transducer operated at a center frequency of 1.5 MHz with an outer active diameter of 25 mm and a 20 ± 2 mm radius of curvature (F# 0.8). The peak negative pressure output of the transducer and its beam profile were calibrated using a needle hydrophone (Onda, Sunnyvale, CA). Acoustic emissions were monitored using a 2 MHz passive cavitation detector (Imasonic, Pessac, France) concentric with the transducer. The detector has a center frequency of 2.02 MHz and a −6 dB fractional bandwidth of 56%, corresponding to a bandwidth of 1.12 MHz (1.46–2.58 MHz). The pulse-echo response has pulse durations of 829 ns at −6 dB and 2.09 μs at −20 dB. This yielded acoustic spectra, from which we extracted quantitative data as the area under the curve of the subharmonic spectral peaks. MBs were infused with a syringe pump (Model Fusion 101A, Chemyx, Texas) via a previously placed custom-made rat catheter (Baxter microbore extension set 2C9201, SAI Infusion Technologies B23-50) at a rate of 50 µL/minute. The concentration of MBs ranged from 20 to 100 µL/kg body weight/minute, diluted in normal saline supplemented with 10% heparin. Animals were anesthetized using less than 3% isoflurane delivered in medical-grade oxygen. The dorsal surface of the skull was shaved to facilitate consistent acoustic coupling between the skin surface and the ultrasound transducer. After the animals were placed in a prone position and the head stabilized, we proceeded with positioning of the FUS target grid. The target grid consisted of subspots separated by 1 mm along the x- and y-directions for a 4 mm by 5 mm grid over the right hemisphere of the animal. MB infusion was allowed to run for 60 sec before sonication commenced. MB-FUS was applied at a duty cycle of ∼2.9-3.1%, a pulse repetition frequency of 1 Hz, with a pulse length of 10 ms per subspot (90 ms OFF period) and sonication within the target grid. Different treatment intensities were achieved by using varying MB dose and system power deposited. The system mechanically steered the acoustic beam over the target grid from one subspot to the next in a recycling pattern. Coupling between the transducer and skin was achieved using mineral oil.

### Mechanical index calculation

Acoustic exposure was quantified as the estimated *in situ* mechanical index (MI). Hydrophone calibration measurements were used to derive the applied acoustic exposure levels. To account for transcranial transmission in rats, we calculated an attenuation coefficient based on body weight, frequency, and skull position in rat-skull transmission data reported by Gerstenmayer et al. and applied this correction to the present study (39). For the mean animal body weight of 280.1 g and a center frequency of 1.5 MHz, the average of skull-position equations yielded a transmission factor of 0.59. The resulting estimated *in situ* mechanical indices were 1.07, 1.60, and 2.13 across the tested exposure conditions.

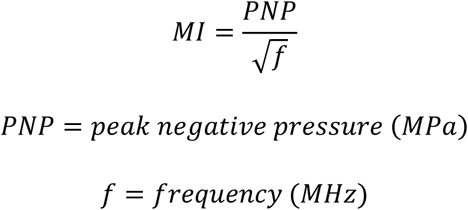

### Acoustic emission dose analysis

Acoustic emission data were analyzed using a standardized dose workflow designed to align preclinical MB-FUS calculations with clinically reported AED (1, 2). For each sonication dataset, paired no-microbubble baseline and microbubble-exposure passive cavitation data (PCD) recordings were analyzed. Recordings were analyzed across 11 MB-FUS treatments, with each treatment containing 20 subspots delivered over 25 cycles. This produced 500 subspot-level AED measurements for each treatment dataset. For each dataset, 100 no-MB baseline sonications and the 500 MB-FUS exposure sonications were analyzed. The primary acoustic emission endpoint was subharmonic AED (AED_SH_) calculated at f0/2 and reported in arbitrary units (a.u.) after baseline normalization. Ultraharmonic AED (AED_UH_) was calculated separately and summed across available ultraharmonic bands. Acquisition metadata, including sampling frequency, shot count, shot duration, fast Fourier transform (FFT) size, voltage conversion factor, and FUS drive frequency, were parsed from the corresponding PCM JSON metadata files. Raw PCD files were read as signed 16-bit integer waveforms and converted to voltage using the metadata-derived voltage conversion factor.

For each pulse, the power spectral density of the voltage waveform was estimated using Welch’s method with a Hann window, the metadata-derived FFT size, and 50% window overlap. Band-integrated acoustic emission power was then calculated by integrating the voltage power spectral density over pre-specified frequency bands centered on the subharmonic frequency, f0/2, and the ultraharmonic frequencies, 3f0/2, 5f0/2, and 7f0/2, where f0 denotes the FUS drive frequency (i.e., fundamental). Together, AED_SH_ and AED_UH_ provide complementary readouts of MB activity during MB-FUS treatments. Subharmonic emissions reflect more nonlinear, potentially unstable oscillations, whereas ultraharmonic emissions provide an additional marker of sustained nonlinear cavitation activity that can help refine acoustic feedback and dosing.

For each emission band *b*, the no-microbubble baseline acoustic emission power, *B_b_*, was calculated as the mean band-integrated emission power across all baseline shots. For each microbubble-exposure shot *j*, a baseline-subtracted acoustic emission score was calculated as follows:

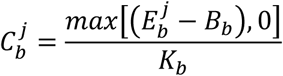

where,

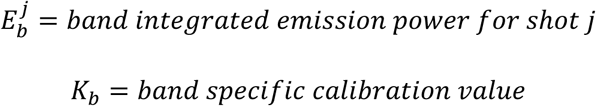

In the primary implementation, K_b_ was set to B_b_, producing a dimensionless, baseline-normalized acoustic emission score reported in arbitrary units. Negative baseline-subtracted values were set to zero before accumulation.

AED_SH_ was pre-specified as the primary AED endpoint because this band most closely follows the clinical AED approach used for MB-FUS BBB opening. For each subspot *s*, AED_SH_ was calculated as follows:

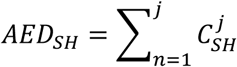

across all sonications delivered to that subspot.

The target-level AED_SH_ was defined as the mean final AED_SH_ across subspots, thereby preserving comparability with clinical target-level AED reporting. The total accumulated AED_SH_ across all shots was also retained as an audit metric but was not used as the primary target-level dose.

AED_UH_ was calculated using the same baseline subtraction, normalization, and accumulation procedure at 3f0/2, 5f0/2, and 7f0/2. Each UH band was normalized to its own no-microbubble baseline before accumulation. A summed AED_UH_ metric was then calculated as the sum of the normalized UH scores across available ultraharmonic bands. Raw ultraharmonic band powers and band-specific AED_UH_ values were exported separately to preserve the contribution of each spectral component.

The broadband inertial cavitation was calculated as a secondary safety control metric. Broadband power was integrated from 1.2f0 to the lower of 2.8f0 or 90% of the usable maximum frequency, while excluding narrow bands around the subharmonic, fundamental, ultraharmonic, and harmonic peaks. Broadband activity was reported in dB above the no-microbubble baseline (***Supplementary Fig. S11***).

For spatial dose mapping, the per-shot acoustic emission scores were organized according to the prescribed subspot-by-cycle firing order. Final per-subspot AED_SH_ and AED_UH_ values were mapped onto the sonication grid using the recorded grid geometry. Both discrete subspot maps and interpolated maps were generated. Interpolation was used only for visualization, while all quantitative analyses were performed on the discrete subspot-level dose values.

### Magnetic resonance imaging

MR imaging was performed on a Bruker 9.4 Tesla Biospec 94/30 USR MR scanner equipped with a range of 1H and multilinear RF coils and interfaced with ParaVision 6.0.1 (Bruker, Ettlingen, Germany). Rats were anesthetized with ∼3% isoflurane in medical-grade oxygen and positioned in an MRI cradle with continuous monitoring of respiratory rate and body temperature (ERT Control/Gating Module, model 1030, SA Instruments). T2-weighted (T2w) imaging was acquired using a rapid spin-echo (RARE) sequence with the following parameters: TR = 2500 ms, TE = 33 ms, RARE factor = 8, FOV = 20×20 mm, matrix = 256×256, slice thickness = 0.5 mm. T1-weighted (T1w) imaging was acquired with TR = 200 ms, TE = 4 ms, FOV = 20×20 mm, matrix = 256×256, slice thickness = 0.5 mm. T1 contrast-enhanced (T1c) imaging was conducted following intravenous administration of Gd-DTPA (0.3 mmol/kg, i.v.; Gadavist®, Bayer, Leverkusen, Germany) using the same T1w parameters. Animals undergoing T1c imaging at later timepoints were given two times the dose of Gd-DTPA intraperitoneally. Images were analyzed to confirm BBB opening and assess any potential off-target effects. Susceptibility-weighted imaging (SWI) was acquired with TR = 350 ms, TE = 16 ms, FOV = 15 mm x 15 mm, matrix = 256 x 256 mm, and slice thickness = 1.0 mm.

### MR image analysis

BBB opening was validated by T1w contrast enhancement, which was analyzed with a custom script in MATLAB R2022b (MathWorks, Natick, MA), based on published methods (3, 40). The analysis began by defining the control ROI (e.g., a non-treated brain region) and the treatment ROI (the FUS-treated and contrast-enhancing brain region). A voxel in the treatment ROI was considered contrast-enhancing if its intensity exceeded the mean intensity of the control ROI by more than two standard deviations. Following this, the BBB opening assessment was normalized to the entire ROI area for each MR image slice. Assessment of hemorrhage and edema was performed using ImageJ (41) to compare pixel intensities within individual MR images. Thresholds for hemorrhage (SWI sequence) and edema (T2w sequence) were empirically derived and determined globally based on visually identified areas of respective findings. The same threshold was applied to all analyzed images. This enabled automatic detection over multiple slices in comparison to the non-sonicated contralateral hemisphere. Each image was verified individually. One image was excluded based on visual assessment of image quality and known MRI sequence artifacts. An additional assessment was performed in which brain slices were segmented by treatment depth: cortex (ctx), ventral subcortex (vsc), dorsal subcortex (dsc), and hippocampus (hc). Differences in BBB opening, microhemorrhage and edema across treatment groups were assessed using one-way ANOVA with Tukey’s HSD post hoc test to correct for multiple comparisons. For treatment depth assessment, differences between groups and treatment intensities were evaluated using two-way ANOVA, with post hoc analyses to control for multiple comparisons. All analyses were completed in GraphPad Prism version 11.

### Clinical trials

The number of relevant clinical trials of MB-FUS for BBB opening was determined by searching ClinicalTrials.gov using the following search parameters: “focused ultrasound” and “blood-brain barrier”. Trials were manually filtered to ensure they were in relation to BBB opening in a relevant indication. Indications included Glioma of all grades and other brain tumors, Alzheimer’s disease, Parkinson’s disease, and Amyotrophic Lateral Sclerosis, among others.

### Behavioral training

Rats were trained on a lever-pressing task using a fully automated home-cage training system, as previously described (27, 42). In short, after initial acclimatization and pre-training, water-restricted rats were rewarded with water for pressing a lever twice within performance-adjusted boundaries around a fixed inter-press interval (IPI) target of 700 ms. Reward boundaries were initially set to 200 ms and 1100 ms. To shape the animals’ behavior toward the target interval while maintaining their motivation to perform the task, the boundaries were dynamically and automatically adjusted between sessions, tightening when the reward rate exceeded 40% and expanding when it was less than 30%. To further incentivize IPIs close to the target, a linearly scaled reward landscape with five evenly spaced levels was established between the target (highest reward, 100 µl water) and each boundary (lowest reward, 20 µL water). Intervals outside the boundaries were not rewarded and triggered a time-out in which animals had to withhold pressing for 1.2 s before initiating a new trial (Inter-trial interval; ITI). Over weeks of training, rats develop increasingly complex and stereotyped kinematics between lever presses (see ‘Kinematic tracking’ below), allowing them to maintain the prescribed IPI target with increasing precision. MB-FUS treatments were performed in expert rats that had reached our previously established learning criterion (27) (mean IPI = 700 ms +/− 10% and coefficient of variation (CV) of IPI distribution < 0.25 in a moving window of 3,000 consecutive trials; typically reached after about 15,000 trials (27); see “Behavioral data analysis” section below), indicating that they had learned the task structure and stabilized their performance.

### Kinematic tracking

Over the course of training, rats develop increasingly complex, spatio-temporally precise, and stereotyped *de novo* movement patterns which span the interval between the lever presses as solutions to our task (27, 29, 30) (cf. ***Fig. 6A***). Raspberry Pi cameras mounted on both sides around the lever captured high-speed videos (90 Hz) of all lever-press related movements throughout training. The recorded videos were automatically cut into 2-second clips, aligned to the first lever press in each trial of the task. To use changes in the kinematics of the developed movement patterns as a behavioral readout for MB-FUS treatment effects, we took advantage of machine learning-aided pose estimation, as in our previous work (29, 30). In short, we used the pose estimation software SLEAP (28) to train a deep neural network on the position of multiple body parts on ∼200 video frames for each trained rat (sampled from before and after treatment). The trained network was then used to predict body-part positions across all frames of all trial videos for the particular rat. We used a Kalman filter to smooth the resulting trajectories and to linearly interpolate the positions of occluded body parts for up to 5 consecutive frames. Trajectories with more than 5 consecutive occluded frames were excluded from analysis.

### Behavioral data analysis

#### Performance metrics

All behavioral analyses were performed using MATLAB 2024a (MathWorks). Performance metrics were determined from lever-press timing in our task. The IPI is the time between the first and second press in a trial, and the ITI is the time between the last press in a non-rewarded trial and the next lever-press. The fraction of trials close to target (trials in the IPI range of 700 ms ± 20%), the IPI, and the CV of the IPI before and after MB-FUS treatment (***Fig. 3, Supplementary Fig. S2***) were calculated over a moving window (25 trials), and the moving average was low-pass filtered with a 50-trial boxcar filter.

#### Criterion performance

We considered animals as experts in our task when their CV was less than 0.25 and the mean of their IPI distribution was in a range of 700 ms ± 10% for a 3,000-trial sliding window. These criteria were previously established and capture the learning performance of intact animals in our task (27).

#### Jensen-Shannon divergence

As a measure for the dissimilarity between interval distributions – either between the IPIs before and after treatment or between the IPIs and ITIs (both before and after treatment) in individual animals, we calculated the Jensen-Shannon divergence (JSD) of the respective contributions. The JSD is a symmetric derivative of the Kullback-Leibler divergence (KLD). For the comparisons of IPIs and ITIs, we calculated the JSD as:

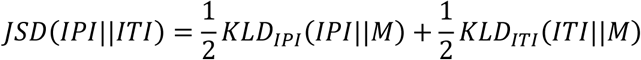

Where

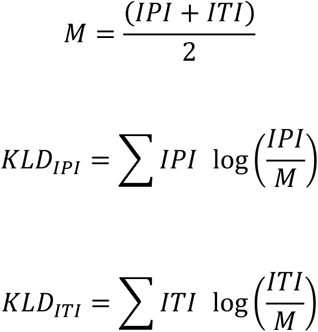

#### Movement trajectory analysis

To determine the effects of MB-FUS treatment on the movement kinematics, we compared the trajectories of both hands of all tracked animals (see Kinematic tracking above) before treatment and on Days 1 and 15 after treatment (***Fig. 6, Supplementary Fig. S7***). Since the hand movements in our task are more pronounced in the vertical than in the horizontal dimension, we focused our analysis on the former. To compare the stereotypy of learned movement patterns, we sub-selected rewarded trials that occurred after unrewarded trials, allowing us to compare trials with the same start and end positions. This is necessary because the rats move up and down between the lever and the water reward port below it after successful trials. Some rats develop more than one movement pattern as solutions to the task, but while these ‘modes’ occur at different frequencies, they all become stereotyped and have similar properties (for details and mode identification see (29, 30)). Here, we sub-selected trials from the most common, dominant mode, so that trajectories are comparable. For illustration purposes, we plotted the average of the selected trajectories with the corresponding Standard Error of the Mean (SEM) of representative example rats (***Fig. 6A***). To calculate the correlations between individual trials (either across all trials on the same day (***Fig. 6B***) or across all trials from a given day and the trials before treatment (***Fig. 6C***)), we linearly warped the trajectories to the same duration (700 ms) by interpolating between the lever-presses. Because the lever-presses themselves have stereotyped trajectories, largely independent of the trial duration, we interpolated the trajectories only from 100 ms after the first to 100 ms before the second lever-press to preserve the shape of the lever-press movements. From these warped trajectories we calculated trial-to-trial correlations separately for both hands and averaged the correlations for each trial. These correlations were averaged for the individual conditions within animals, and those means were plotted, together with violin plots and boxplots (***Fig. 6B, C***). We also compared the distributions of these different correlations for individual animals (examples, see ***Fig. 6D***): we compared the distributions of correlations for trials before treatment (before vs. before) to the distributions of correlations across before treatment and Day 1 or Day 15 and used a two-sided KS-test to determine for individual animals whether their before:before distributions were significantly different from their before:Day 1 or before:Day 15 distributions (***Fig. 6E***), indicating behavioral changes.

#### Statistics

The significance of observed acute effects of the different MB-FUS treatments on behavioral measures was tested using One-way ANOVAs with treatment type as a fixed factor. When significant differences were detected, post hoc tests with the Holm correction for multiple comparisons were performed. For comparisons of MB-FUS treatment effects across days (repeated measures), we fitted linear mixed-effect models, using treatment type and days as fixed factors and random intercepts varying by animal to account for individual variability. The models followed the template: (outcome ∼ Treatment * Days + (1|Rat)). Subsequently, we estimated marginal means and planned pairwise contrasts for each day between treatments and for each treatment across days (holding the other factor constant), with family-wise adjustments using the ‘mvt’ method (multivariate t adjustment). We used the statistics software JASP (0.95.4, 2025, University of Amsterdam) for statistical analysis. For ANOVAs, we used the built-in functionality. For linear mixed effect models and contrast testing, we provided custom commands via JASP’s R console, using the ‘lmer’ (‘lme4’ R package) and ‘emmeans’ (‘emmeans’ R package) functions. The KS-test to determine whether correlation distributions are similar or distinct (***Fig. 6E***) was performed in Matlab 2024A (Mathworks). Significant differences are indicated in the respective figures, and statistical details for behavioral analyses are provided in supplementary tables associated with individual figures.

### Histology and histological analysis

Naïve, untrained rats (n=15) received the four highest MB-FUS doses (Ultra-High/D, High/D, High/S, and Medium/D; see ***Table 1***). 3 days after MB-FUS treatment, rats were given inhalation anesthesia (isoflurane) and perfused with phosphate-buffered saline (PBS), followed by 4% paraformaldehyde (PFA) in PBS. Their brains were harvested for histological analysis, sectioned into 80 μm coronal slices on a vibratome (Leica VT1000S), and underwent immunofluorescence staining to determine the extent and location of MB-FUS-induced astrocyte activation. Sections were blocked for 2 hours on a rocker at room temperature in blocking solution (1% bovine serum albumin and 0.3% Triton-X 100 in PBS) and stained overnight at room temperature with a primary antibody for glial fibrillary acidic protein (GFAP, to stain for astrocytes; 1:500 in blocking solution; Sigma Aldrich, G9269, raised in rabbit). After subsequent washing (3x for 10 min with PBS), sections were stained with fluorescently labeled secondary antibodies (1:1000 in blocking solution; anti-rabbit, Alexa Fluor 488; Invitrogen, A11008) for 2 hours at room temperature. Slides were subsequently washed (3x, 10 min each) in PBS and mounted with Fluoromount-G mounting medium containing DAPI (Invitrogen, 00-4959-52) to visualize cell nuclei. Images of whole brain slices were acquired at 4x magnification with a BZ-X710 All-in-One Fluorescence Microscope (Keyence).

To quantify the difference in GFAP intensity between treated and untreated hemispheres, ROIs for multiple brain areas were manually drawn in ImageJ (cortex, hippocampus, dorsal subcortex, and ventral subcortex; see ***Supplementary Fig. S4A***), and their mean fluorescence intensity was measured. Using custom Python code, we calculated the intensity ratio between the treated and untreated hemispheres for each ROI across six representative planes (+3.20 mm, +1.60 mm, +0.70 mm, −0.40 mm, −2.30 mm, and –3.80 mm from Bregma).

## Data, Materials and Software availability

Raw data, including MRI and histological images and behavioral data, as well as analysis code, will be made publicly available upon publication of this manuscript.

## Acknowledgements

This work was supported by a National Institute of Mental Health grant R01 MH136968, and a fellowship from the Esther A. & Joseph Klingenstein Fund and the Simons Foundation, both awarded to S.B.E.W., an American Cancer Society’s Institutional Research Grant IRG-18-160-16 to P.A., and a National Cancer Institute of the National Institutes of Health grant T32CA154274 awarded to A.A.S. The content is solely the responsibility of the authors and does not necessarily represent the official views of the National Institutes of Health. We thank members of the Wolff and Anastasiadis labs for technical support and feedback. We also thank Hendrik Wolff, Tim Faw, and Erik Dumont for discussions and feedback to the manuscript, and biological illustrator Tina Wang for her contributions to Figures 1 and 3A.

## Author Contributions

S.K.E. and A.A.S. contributed equally. S.B.E.W. and P.A. are co-corresponding authors. S.K.E., A.A.S., P.A. and S.B.E.W. designed the study, performed experiments, analyzed the data, and wrote the manuscript.

## Competing interests

The authors declare no competing interests.

## Notes

### Competing Interest Statement

The authors have declared no competing interest.

