## Supplementary Figure S1 for "Neurobehavioral effects of focused ultrasound-mediated blood-brain barrier opening"

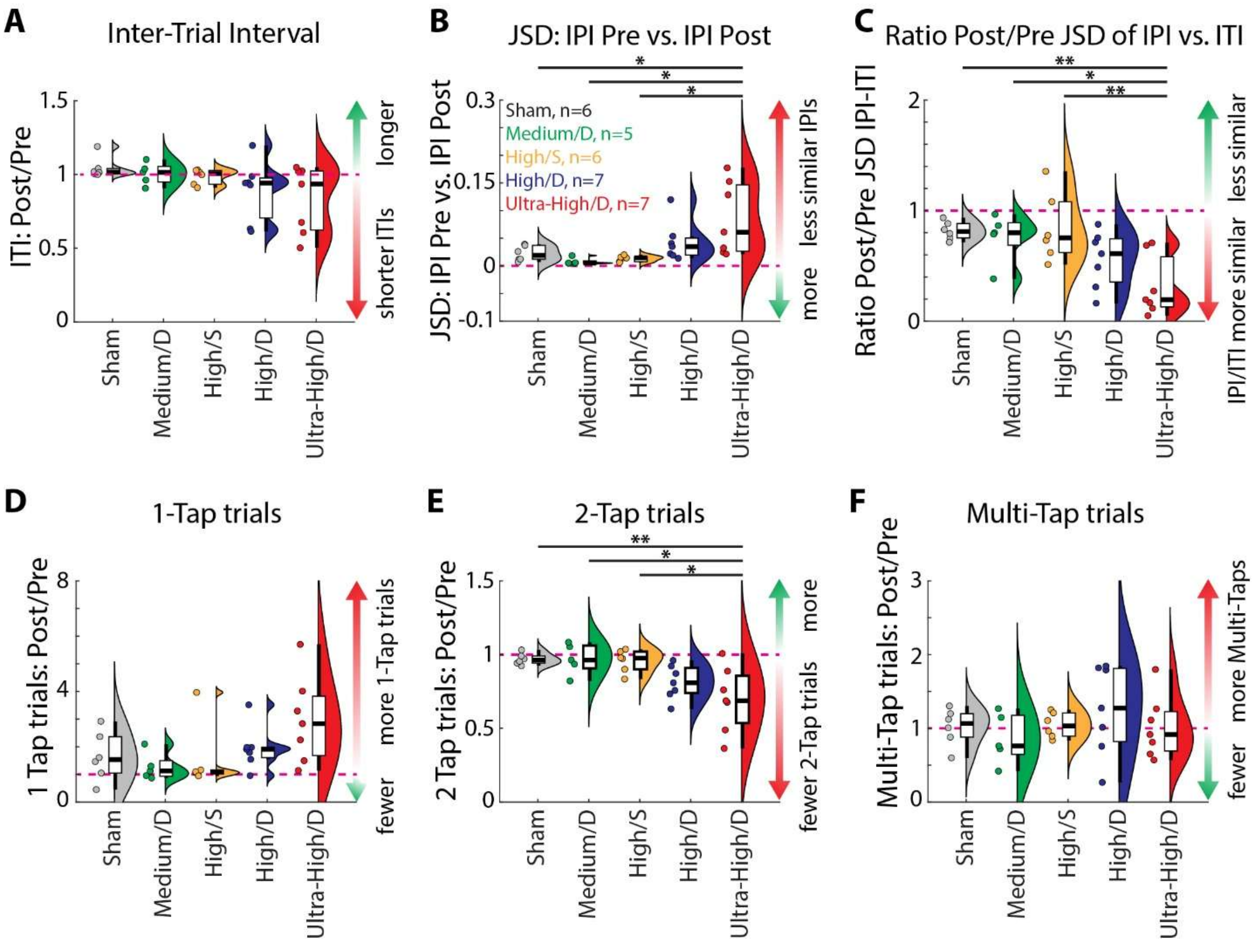


**Supplementary Figure S1. Additional acute performance measures for high-intensity MB-FUS treatment regimens. A** Ratios of the Inter-Trial intervals (ITI) between post- (1,000 trials) and pre- (2,000 trials) MB-FUS treatment. Rats need to keep a minimum ITI of 1.2 s after unrewarded trials before initiating a new trial. See Methods for details. **B** Jenson-Shannon Divergence (JSD) between pre- and post-treatment IPIs as a measure for the dissimilarity of the IPI distributions. Higher JSD values indicate a higher dissimilarity between the pre- and post- IPI distributions. **C** Ratios between the post- and pretreatment JSD values between the IPI and ITI distributions. Rats learn to produce distinct IPIs and ITIs, reflected by an increasing JSD values across learning. Comparing these values across post- and pre-treatment indicates whether the animals' ability to generate distinct intervals is impaired by the treatment. Post-/Pre- ratios below 1 indicate that animals increased similarity between IPIs and It is, i.e., less distinction. **D** Ratios of the fraction of trials with only 1 lever press between post-and pre- MB-FUS treatment. Only trials with exactly 2 lever presses can be rewarded. **E** Like D, but for trials with exactly 2 lever presses. **F** Like D, but for trials with more than 2lever presses. Statistics: 1-way ANOVAs with Post-hoc tests (Holm), see ***Supplementary Table S3*** for detailed results. Significance levels: * $p<0.05$; ** $p<0.01$; ***$p<0.001$.
