## Supplementary Figure S2 for "Neurobehavioral effects of focused ultrasound-mediated blood-brain barrier opening"

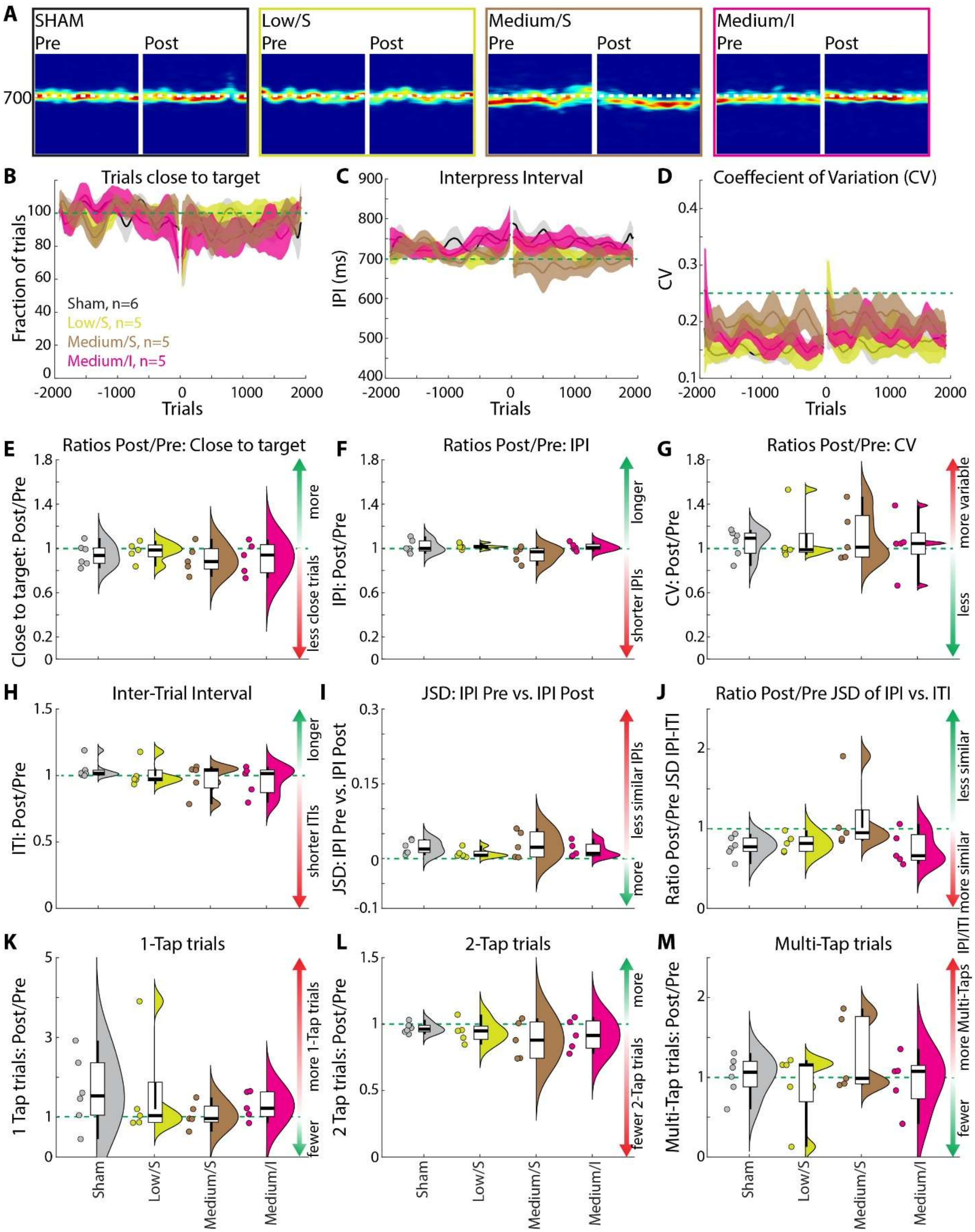


**Supplementary Figure S2. Acute performance measures for low-intensity MB-FUS treatment regimens. A** Example performance plots of rats treated with distinct MB-FUS regimens. Probability density plots show the likelihood of the generation of trials with specific IPIs over time, for 1,000 trials pre- and post MB-FUS treatment. Warmer colors indicate higher likelihoods. **B** Population plot for the fraction of trials close to the target pre- and post-MB-FUS treatment for multiple MB-FUS regimens and sham-treated rats, normalized to pre-treatment performance. Shown are population means and SEM. **C** Population plot for the inter-press interval pre- and post-MB-FUS treatment. **D** Population plot of the CV of the IPI as a measure of behavioral variability pre- and post-MB-FUS treatment. **E** Ratios of the fraction of trials close to the target between post- (1,000 trials) and pre- (2,000 trials) MB-FUS treatment. Violin plots with boxplots and individual rats as in ***Fig. 3G***. **F** As E, but for the ratios of IPIs. **G** As E, but for the ratios of CVs. **H** Ratios of the Inter-Trial intervals (ITI) between post- (1,000 trials) and pre- (2,000 trials) MB-FUS treatment. **I** Jenson-Shannon Divergence (JSD) between pre- and post-treatment IPIs as a measure for the dissimilarity of the IPI distributions. **J** Ratios between the post- and pretreatment JSD values between the IPI and ITI distributions. **K** Ratios of the fraction of trials with only 1 lever press between post- and pre- MB-FUS treatment. Only trials with exactly 2 lever presses can be rewarded. **L** Like K, but for trials with exactly 2 lever presses. **M** Like K, but for trials with more than 2 lever presses. Statistics (E-M): 1-way ANOVAs with Post-hoc tests (Holm), see ***Supplementary Table S4*** for detailed results.
