## Supplementary Figure S3 for "Neurobehavioral effects of focused ultrasound-mediated blood-brain barrier opening"

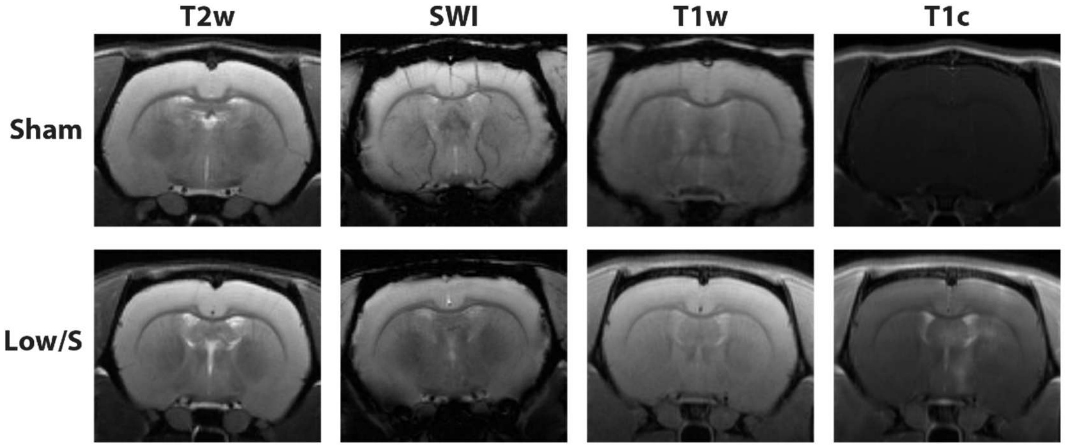


**Supplementary Figure S3. Effects of sham and lowest dose MB-FUS treatments.** MRI sequences from example rats from the sham (top) and Low/S (bottom) treatment groups, taken within 30 minutes after treatment. The T1c sequence indicates successful BBB opening in the Low/S treatment rat, but no BBB opening in the sham-treated rat. Additional MRI sequences do not show any tissue damage in either example rat.
