## Supplementary Figure S4 for "Neurobehavioral effects of focused ultrasound-mediated blood-brain barrier opening"

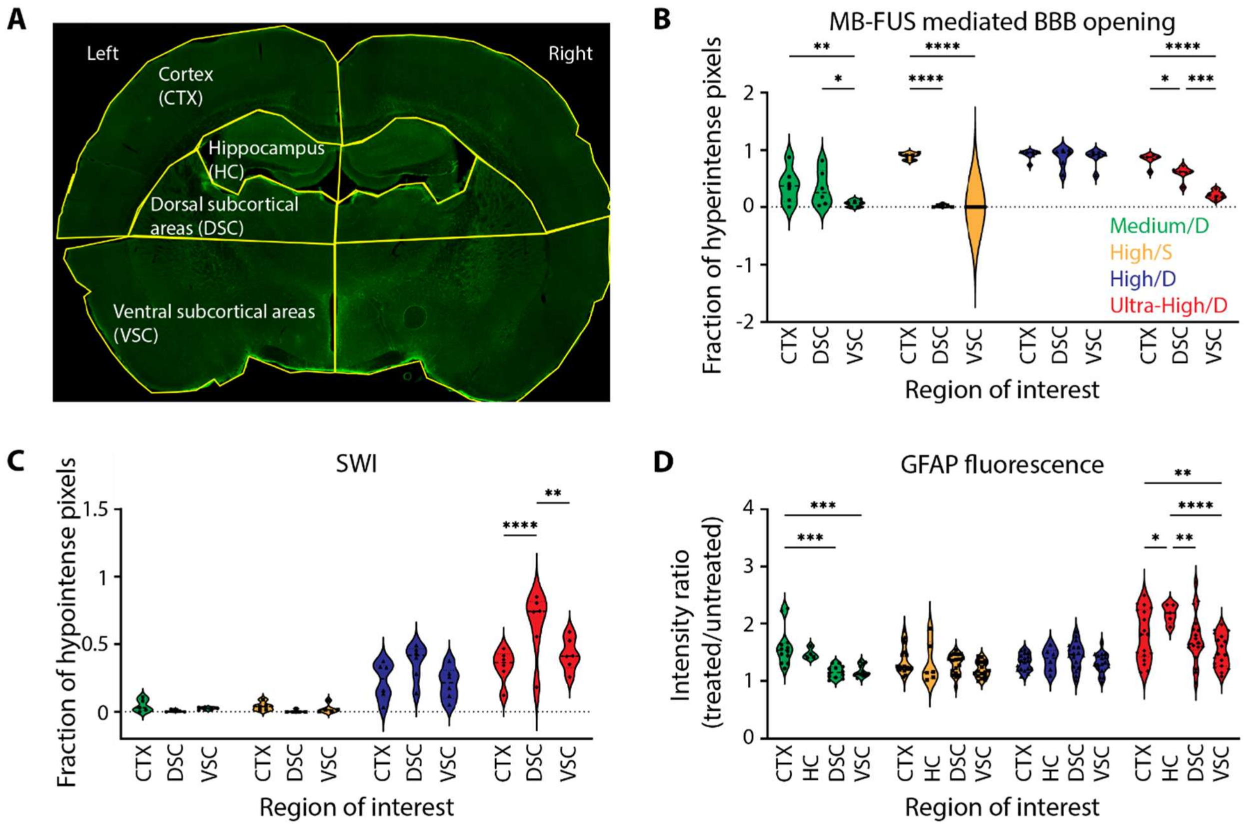


**Supplementary Figure S4. Quantification of MRI sequences and histological assessment by depth. A** Example of ROIs for quantifications of MB-FUS induced effects: cortex (CTX), hippocampus (HC), dorsal (DSC), and ventral subcortical (VSC) areas. Note that the subcortical ROIs are determined only based on depth, not based on anatomically defined brain areas. **B** Population quantifications of T1w MRI sequence after MB-FUS treatments from ***Fig. 4*** shown by ROI. **C** Like B, but for the SWI MRI sequence. **D** Like B, but for GFAP fluorescence. Statistics: Only comparisons across depths in the same treatment group are shown, for all other pairwise comparisons, see ***Supplementary Table S6***. 2-way ANOVA with multiple comparisons corrections. * $p<0.05$; ** $p<0.01$; *** $p<0.001$; **** $p<0.0001$.
