## Supplementary Figure S5 for "Neurobehavioral effects of focused ultrasound-mediated blood-brain barrier opening"

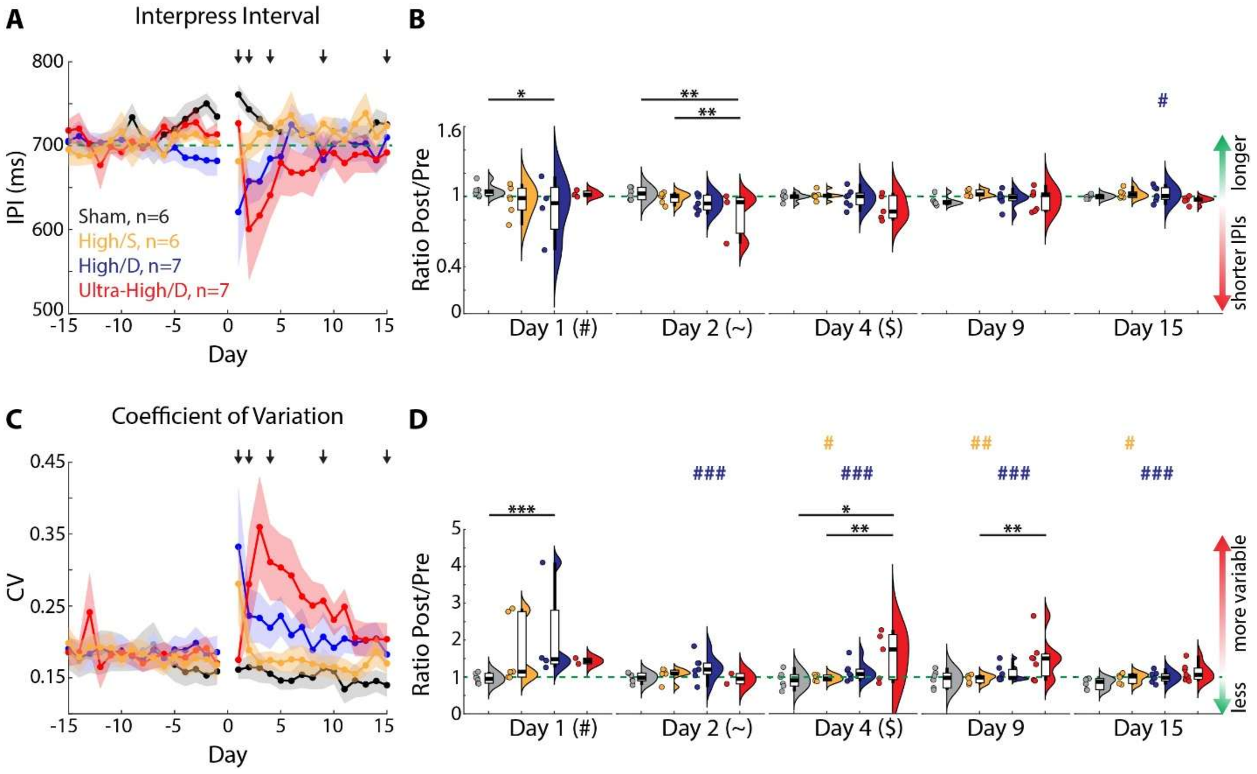


**Supplementary Figure S5. Additional long-term performance measures for high-intensity MB-FUS treatment regimens. A** IPIs across days pre- and post-MB-FUS treatment. IPIs are averaged across two daily sessions, excluding missed sessions without trials. Dots: Population average per day. Error bars: SEM. **B** Ratios of IPIs between post- and pre-treatment (averaged across 10 days) for selected days. Only rats with active sessions on a given day are included. **C, D** Like A,B but for the CV. Violin plots with boxplots and individual rats as in ***Fig. 3G***. Statistics (B,D): Linear Mixed Model with planned contrasts (Comparison across groups for each day and comparisons across days for each group). For detailed results see ***Supplementary Table S8***. Stars indicate significant differences across groups on the same day. Colored symbols indicate significant differences across days for a given group (Symbol indicates compared day (e.g., # compared to Day 1), color indicates the group (e.g., red for Ultra-High/D group). Significance levels: * p<0.05; ** p<0.01; ***p<0.001.
