## Supplementary Figure S6 for "Neurobehavioral effects of focused ultrasound-mediated blood-brain barrier opening"

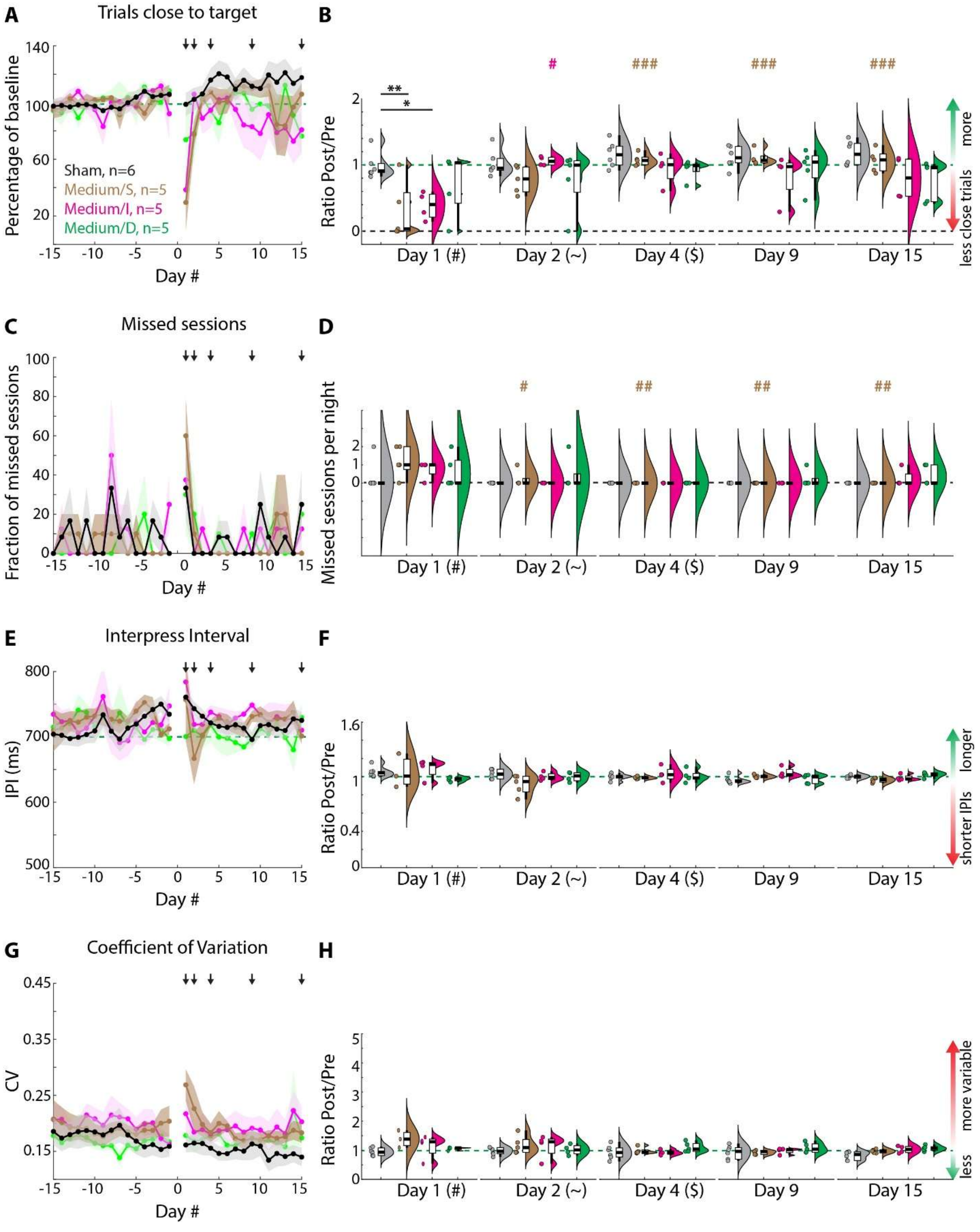


**Supplementary Figure S6. Long-term performance measures for low-intensity MB-FUS treatment regimens. A** Trials close to the target across days pre- and post-MB-FUS treatment, normalized to pre-treatment performance. Daily performance is averaged across two daily sessions. Missed sessions are considered as having a fraction of trials close to target of 0. Dots: Population average per day. Error bars: SEM. **B** Ratios of the fraction of trials close to the target between post- and pre-treatment (averaged across 10 days) for selected days. **C** Like A, but for the fraction of missed sessions per day. Rats can perform in up to two sessions per night. Sessions are counted as missed, if no trials are performed during the session time. **D** Like B, but for the number of missed sessions per night. **E,F** Like A,B but for the CV. **G, H** Like A,B but for the CV. Violin plots with boxplots and individual rats as in ***Fig. 3G***. Statistics (B,D): Linear Mixed Model with planned contrasts (Comparison across groups for each day and comparisons across days for each group). For detailed results see ***Supplementary Table S9***. Stars indicate significant differences across groups on the same day. Colored symbols indicate significant differences across days for a given group (Symbol indicates compared day (e.g., # compared to Day 1), color indicates the group (e.g., red for Ultra-High/D group). Significance levels: * $p<0.05$; ** $p<0.01$; ***$p<0.001$.
