## Supplementary Figure S7 for "Neurobehavioral effects of focused ultrasound-mediated blood-brain barrier opening"

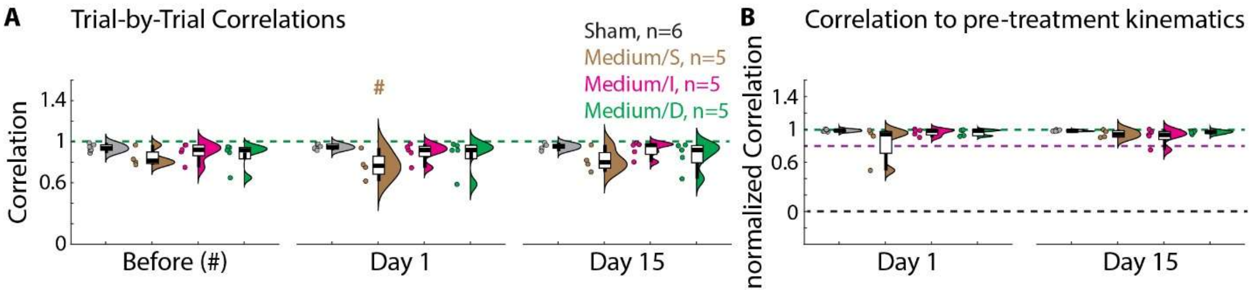


**Supplementary Figure S7. Acute and long-term effects of low-intensity MB-FUS treatment on the complex learned kinematics in a motor skill task. A** Trial-by-trial correlations across all trials before MB-FUS treatment (across 5 days), 1 day or 15 days after treatment. **B** Correlation between trials post- and pre-MB-FUS treatment, normalized to the trial-by-trial correlation pre-treatment for each animal (see B). Violin plots with boxplots and individual rats as in ***Fig. 3G***. Statistics: Linear Mixed Model with planned contrasts (Comparison across groups for each day and comparisons across days for each group). For detailed results see ***Supplementary Table S11***. Stars indicate significant differences across groups on the same day. Colored symbols indicate significant differences across days for a given group (Symbol indicates compared day (e.g., # compared to Day 1), color indicates the group (e.g., brown for Medium/S group). Significance levels: * p<0.05; ** p<0.01; ***p<0.001.
