## Supplementary Figure S8 for "Neurobehavioral effects of focused ultrasound-mediated blood-brain barrier opening"

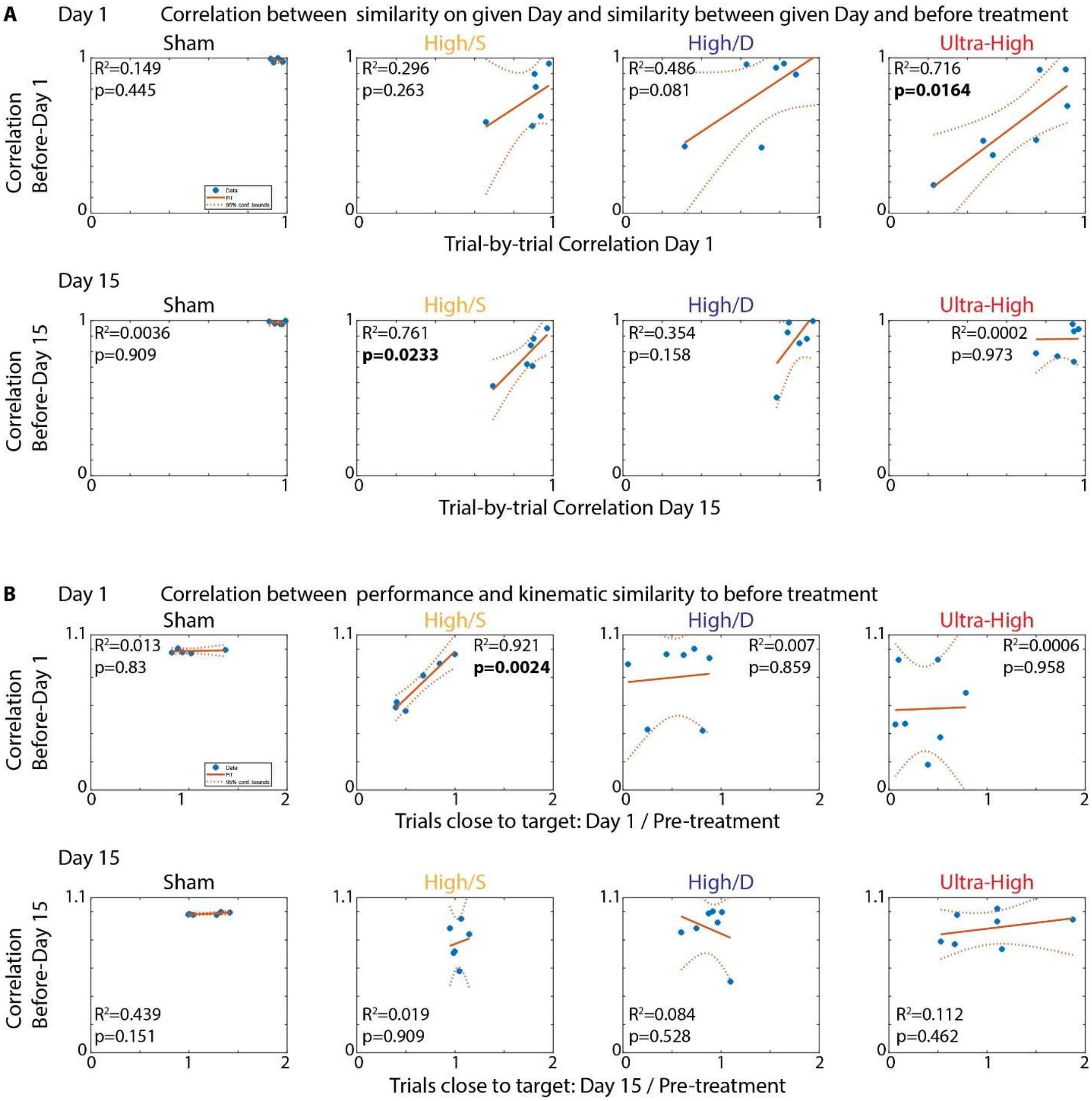


**Supplementary Figure S8. Reduced similarity between before- and after-treatment trajectories cannot be explained by reduced stereotypy alone and does not correlate with reduced performance. A** Correlations between movement variability after treatment and similarity between pre-and-post treatment movements. Shown are the correlations between trial-by-trial correlations on Day 1 (top) and Day 15 (bottom) and the correlations between the trajectories before treatment and the trajectories on Day 1 (or 15). Shown are individual animals (dots), linear trendlines (solid lines) and 95% confidence intervals (dotted lines). **B** As A, but for correlations between performance (Post/Pre ratio for the trials close to the target on the given day; see ***Fig. 5A***) and the correlations between the trajectories before treatment and the trajectories on Day 1 (or 15).
