## Supplementary Figure S9 for "Neurobehavioral effects of focused ultrasound-mediated blood-brain barrier opening"

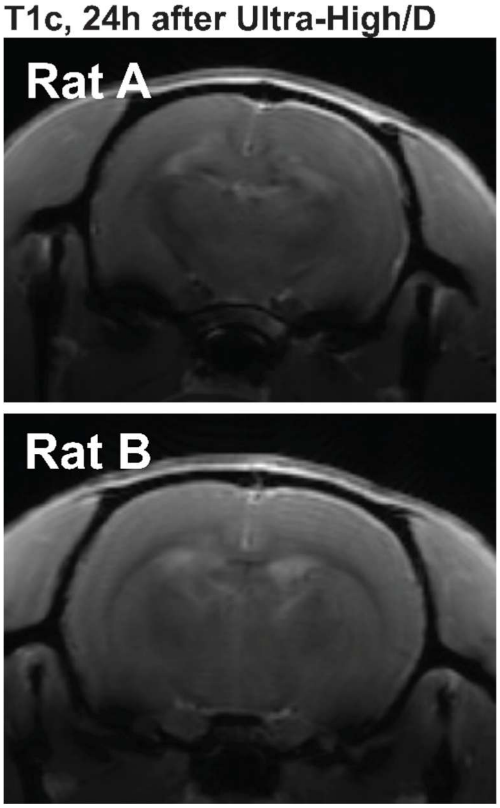


**Supplementary Figure S9: T1c scans 24 hours after MB-FUS treatment.** Example rats from the Ultra-High/D treatment group. 24 hours after treatment, rats received a factor 10 dose of gadolinium and underwent T1w MRI scans. Visual inspection shows no gadolinium contrast enhancement in the treated area, suggesting re-sealing of the BBB.
