## Supplementary Figure S10 for "Neurobehavioral effects of focused ultrasound-mediated blood-brain barrier opening"

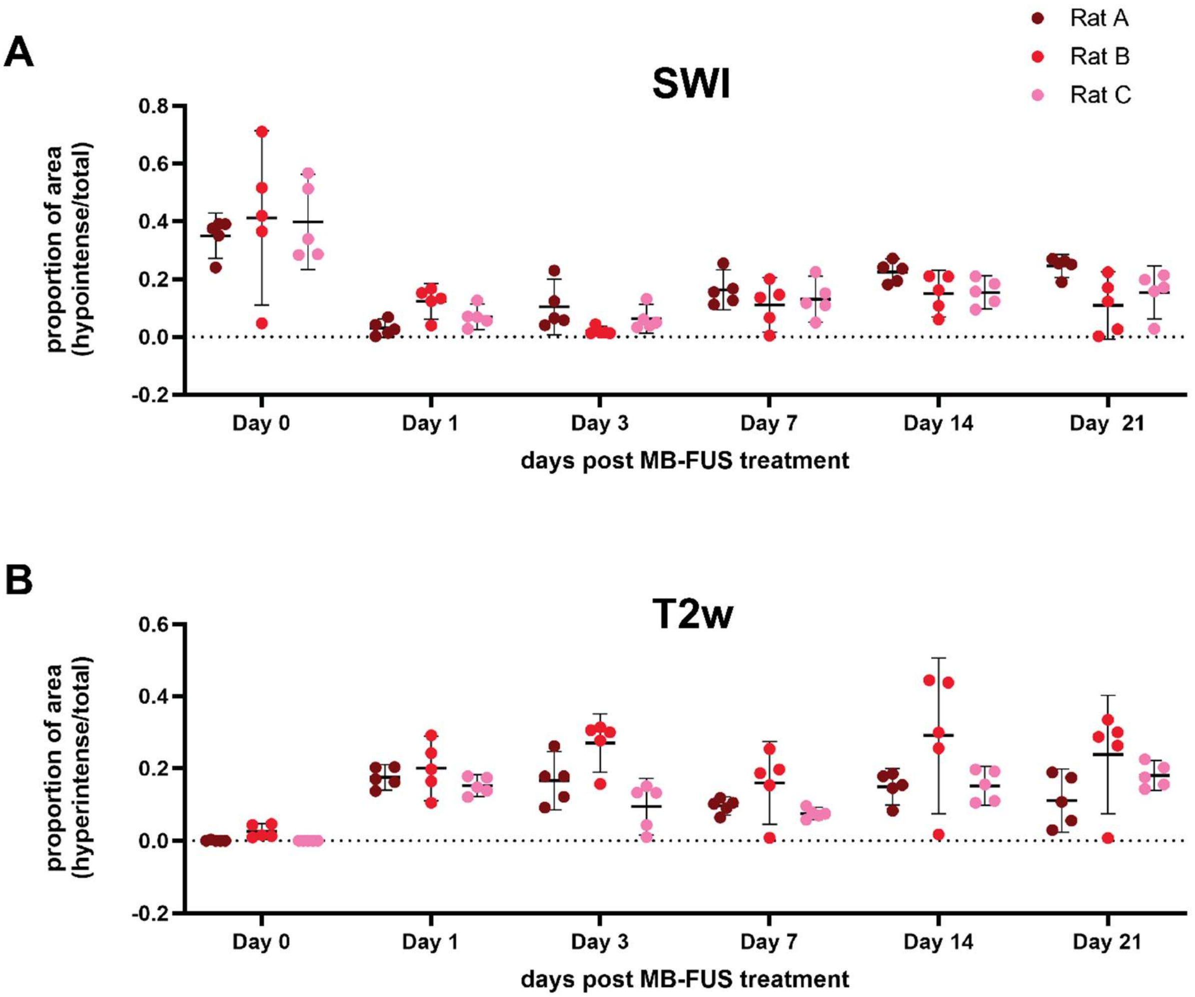


**Supplementary Figure S10. Quantification of longitudinal imaging after Ultra-High/D MB-FUS treatment for individual animals. A** Quantification of the SWI MRI sequence (see ***Fig. 7***) for individual animals. Dots represent the individual imaging planes for each animal. **B** As A, but for the T2w MRI sequence.
