## Supplementary Figure S11 for "Neurobehavioral effects of focused ultrasound-mediated blood-brain barrier opening"

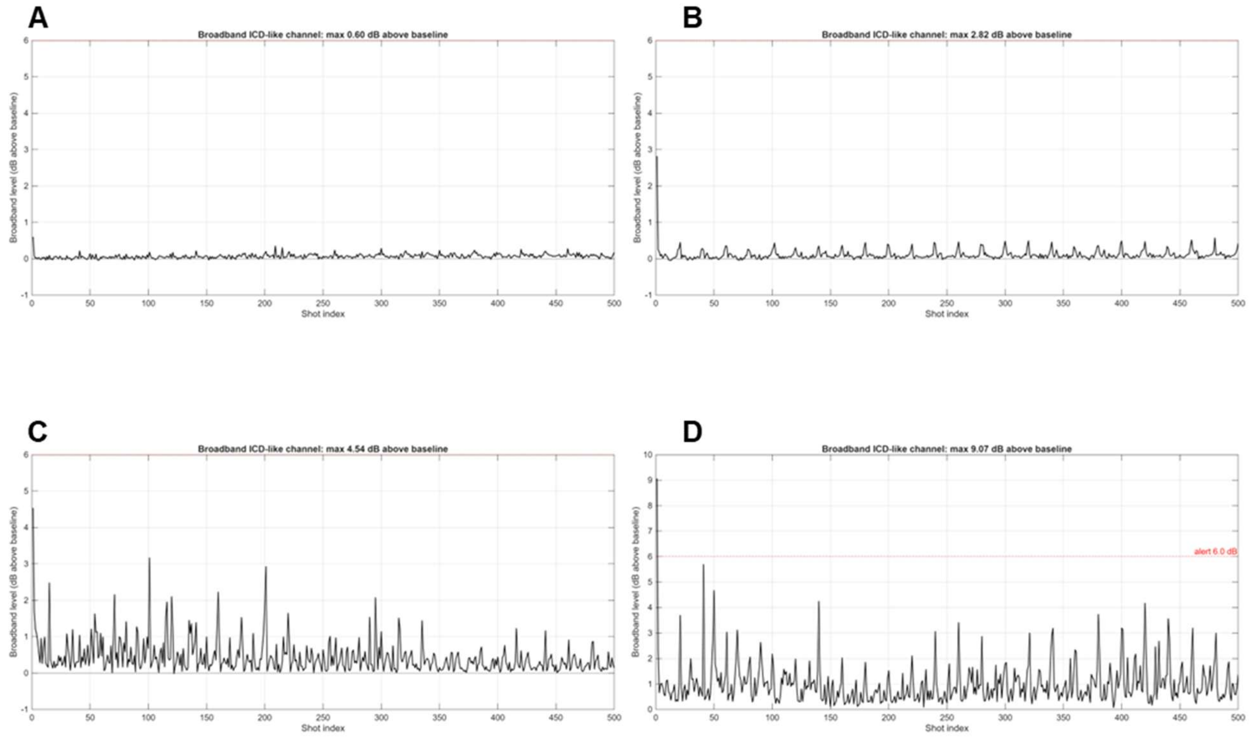


**Supplementary Figure S11. Broadband inertial cavitation for each experimental group.** Emissions are reported in decibels (dB) above baseline over the course of the 500 treatment sonications, indicating unstable collapse of microbubbles at that shot. **A** Representative broadband cavitation for a Medium/D treatment. **B-D** As A, for High/S, High/D, and Ultra-High/D treatments.
