## Supplementary Table S1 for "Neurobehavioral effects of focused ultrasound-mediated blood-brain barrier opening"

**Supplementary Table S1. Median $AED_{SH}$ and $AED_{UH}$ and interquartile range (IQR) for each treatment.** Related to Fig. 2.

| Treatment | Median $AED_{SH}$ | $AED_{SH}$ IQR | Median $AED_{UH}$ | $AED_{UH}$ IQR |
|---|---|---|---|---|
| **Medium/D** | 2.15 | 0.21-5.29 | 1.96 | 0.27-6.33 |
| **High/S** | 12.06 | 2.50-17.18 | 2.70 | 0.02-17.13 |
| **High/D** | 30.67 | 17.90-54.59 | 29.71 | 9.32-83.43 |
| **Ultra-High/D** | 111.00 | 66.12-159.9 | 68.36 | 35.71-129.80 |
