## Supplementary Table S2 for "Neurobehavioral effects of focused ultrasound-mediated blood-brain barrier opening"

### Supplementary Table S2*: Statistical Results for Figure 3*

| Figure | Description | Samples | Test | Results |
|---|---|---|---|---|
| Fig. 3G | Comparison of ratios between post- and pre-treatment values for the fraction of trials close to the target (+/-20% around 700 ms) across treatment groups | Rat numbers:<br>Sham: 6<br>Medium/D: 5<br>High/S: 6<br>High/D: 7<br>Ultra-High/D: 7 | One-Way ANOVA. Factor 1: Treatment | Treatment: F(4,26)=10.80, P<0.001 |
| | | | Post-hoc test: Holm | P values:<br>1)Sham vs. Medium/D: P=1<br>2)Sham vs. High/S: P=1<br>**3)Sham vs. High/D: P=0.03**<br>**4)Sham vs. U-H/D: P<0.001**<br>5)Medium/D vs. High/S: P=1<br>**6)Medium/D vs. High/D: P=0.03**<br>**7)Medium/D vs. U-H/D: P<0.001**<br>**8)High/S vs. High/D: P=0.031**<br>**9)High/S vs. U-H/D: P<0.001**<br>10)High/D vs. U-H/D: P=0.233 |
| Fig. 3H | Comparison of ratios between post- and pre-treatment IPIs across treatment groups | Rat numbers:<br>Sham: 6<br>Medium/D: 5<br>High/S: 6<br>High/D: 7<br>Ultra-High/D: 7 | One-Way ANOVA. Factor 1: Treatment | Treatment: F(4,26)=1.576, P=0.210 |
| Fig. 3I | Comparison of ratios between post- and pre-treatment values for the CV of IPIs across treatment groups | Rat numbers:<br>Sham: 6<br>Medium/D: 5<br>High/S: 6<br>High/D: 7<br>Ultra-High/D: 7 | One-Way ANOVA. Factor 1: Treatment | Treatment: F(4,26)=6.903, P<0.001 |
| | | | Post-hoc test: Holm | P values:<br>1)Sham vs. Medium/D: P=1<br>2)Sham vs. High/S: P=1<br>3)Sham vs. High/D: P=0.473<br>**4)Sham vs. U-H/D: P=0.003**<br>5)Medium/D vs. High/S: P=1<br>6)Medium/D vs. High/D: P=0.288<br>**7)Medium/D vs. U-H/D: P=0.002**<br>8)High/S vs. High/D: P=0.520<br>**9)High/S vs. U-H/D: P=0.005**<br>10)High/D vs. U-H/D: P=0.146 |
