## Supplementary Table S3 for "Neurobehavioral effects of focused ultrasound-mediated blood-brain barrier opening"

**Supplementary Table S3*: Statistical Results for Supplementary Figure S1.***

| Figure | Description | Samples | Test | Results |
|---|---|---|---|---|
| Fig. S1A | Comparison of ratios between post- and pre-treatment ITIs across treatment groups | Rat numbers:<br>Sham: 6<br>Medium/D: 5<br>High/S: 6<br>High/D: 7<br>Ultra-High/D: 7 | One-Way ANOVA. Factor 1: Treatment | Treatment: F(4,26)=2.05, P=0.117 |
| Fig. S1B | Comparison of JSD values for IPI distributions post- and pre-treatment across treatment groups | Sham: 6<br>Medium/D: 5<br>High/S: 6<br>High/D: 7<br>Ultra-High/D: 7 | One-Way ANOVA. Factor 1: Treatment | Treatment: F(4,26)=4.760, P=0.005 |
| | | | Post-hoc test: Holm | P values:<br>1)Sham vs. Medium/D: P=1<br>2)Sham vs. High/S: P=1<br>3)Sham vs. High/D: P=1<br>**4)Sham vs. U-H/D: P=0.038**<br>5)Medium/D vs. High/S: P=1<br>6)Medium/D vs. High/D: P=0.656<br>**7)Medium/D vs. U-H/D: P=0.013**<br>8)High/S vs. High/D: P=0.753<br>**9)High/S vs. U-H/D: P=0.014**<br>10)High/D vs. U-H/D: P=0.294 |
| Fig. S1C | Comparison of ratios between post- and pre-treatment values for the JSD of IPI and ITI distributions across treatment groups | Sham: 6<br>Medium/D: 5<br>High/S: 6<br>High/D: 7<br>Ultra-High/D: 7 | One-Way ANOVA. Factor 1: Treatment | Treatment: F(4,26)=5.614, P=0.002 |
| | | | Post-hoc test: Holm | P values:<br>1)Sham vs. Medium/D: P=1<br>2)Sham vs. High/S: P=1<br>3)Sham vs. High/D: P=0.348<br>**4)Sham vs. U-H/D: P=0.007**<br>5)Medium/D vs. High/S: P=1<br>6)Medium/D vs. High/D: P=0.678<br>**7)Medium/D vs. U-H/D: P=0.034**<br>8)High/S vs. High/D: P=0.307<br>**9)High/S vs. U-H/D: P=0.006**<br>10)High/D vs. U-H/D: P=0.351 |
| Fig. S1D | Comparison of ratios between post- and pre-treatment fractions of 1-tap trials across treatment groups | Sham: 6<br>Medium/D: 5<br>High/S: 6<br>High/D: 7<br>Ultra-High/D: 7 | One-Way ANOVA. Factor 1: Treatment | Treatment: F(4,26)=2.364, P=0.079 |

| | | | | |
|---|---|---|---|---|
| Fig. S1E | Comparison of ratios between post- and pre-treatment fractions of 2-tap trials across treatment groups | Sham: 6<br>Medium/D: 5<br>High/S: 6<br>High/D: 7<br>Ultra-High/D: 7 | One-Way ANOVA. Factor 1: Treatment | Treatment: $F(4,26)=5.641$, $P=0.002$ |
| | | | Post-hoc test: Holm | P values:<br>1)Sham vs. Medium/D: P=1<br>2)Sham vs. High/S: P=1<br>3)Sham vs. High/D: P=2.92<br>**4)Sham vs. U-H/D: P=0.009**<br>5)Medium/D vs. High/S: P=1<br>6)Medium/D vs. High/D: P=0.293<br>**7)Medium/D vs. U-H/D: P=0.013**<br>8)High/S vs. High/D: P=0.295<br>**9)High/S vs. U-H/D: P=0.013**<br>10)High/D vs. U-H/D: P=0.443 |
| Fig. S1F | Comparison of ratios between post- and pre-treatment fractions of Multi-tap trials across treatment groups | Sham: 6<br>Medium/D: 5<br>High/S: 6<br>High/D: 7<br>Ultra-High/D: 7 | One-Way ANOVA. Factor 1: Treatment | Treatment: $F(4,26)=0.728$, $P=0.581$ |
