## Supplementary Table S4 for "Neurobehavioral effects of focused ultrasound-mediated blood-brain barrier opening"

**Supplementary Table S4*: Statistical Results for Supplementary Figure S2.***

| Figure | Description | Samples | Test | Results |
|---|---|---|---|---|
| Fig. S2E | Comparison of ratios between post- and pre-treatment for fraction of trials close to target across treatments | Rat numbers:<br>Sham: 6<br>Low/S: 5<br>Medium/S: 5<br>Medium/I: 5 | One-Way ANOVA. Factor 1: Treatment | Treatment: F(3,17)=0.357, P=0.784 |
| Fig. S2F | Comparison of ratios between post- and pre-treatment IPIs across treatment groups | Sham: 6<br>Low/S: 5<br>Medium/S: 5<br>Medium/I: 5 | One-Way ANOVA. Factor 1: Treatment | Treatment: F(3,17)=2.495, P=0.095 |
| Fig. S2G | Comparison of ratios between post- and pre-treatment values for the CV of IPIs across treatment groups | Sham: 6<br>Low/S: 5<br>Medium/S: 5<br>Medium/I: 5 | One-Way ANOVA. Factor 1: Treatment | Treatment: F(3,17)=0.123, P=0.945 |
| Fig. S2H | Comparison of ratios between post- and pre-treatment ITIs across treatment groups | Sham: 6<br>Low/S: 5<br>Medium/S: 5<br>Medium/I: 5 | One-Way ANOVA. Factor 1: Treatment | Treatment: F(3,17)=0.755, P=0.535 |
| Fig. S2I | Comparison of JSD values for IPI distributions post- and pre-treatment across treatment groups | Sham: 6<br>Low/S: 5<br>Medium/S: 5<br>Medium/I: 5 | One-Way ANOVA. Factor 1: Treatment | Treatment: F(3,17)=0.947, P=0.440 |
| Fig. S2J | Comparison of ratios between post- and pre-treatment values for JSD of IPI and ITI distributions across treatment groups | Sham: 6<br>Low/S: 5<br>Medium/S: 5<br>Medium/I: 5 | One-Way ANOVA. Factor 1: Treatment | Treatment: F(3,17)=1.694, P=0.206 |
| Fig. S2K | Comparison of ratios between post- and pre-treatment fractions of 1-tap trials across treatment groups | Sham: 6<br>Low/S: 5<br>Medium/S: 5<br>Medium/I: 5 | One-Way ANOVA. Factor 1: Treatment | Treatment: F(3,17)=0.572, P=0.641 |
| Fig. S2L | Comparison of ratios between post- and pre-treatment fractions of 2-tap trials across treatment groups | Sham: 6<br>Low/S: 5<br>Medium/S: 5<br>Medium/I: 5 | One-Way ANOVA. Factor 1: Treatment | Treatment: F(3,17)=0.731, P=0.548 |
| Fig. S2L | Comparison of ratios between post- and pre-treatment fractions of Multi-tap trials across treatment groups | Sham: 6<br>Low/S: 5<br>Medium/S: 5<br>Medium/I: 5 | One-Way ANOVA. Factor 1: Treatment | Treatment: F(3,17)=0.934, P=0.446 |
