## Supplementary Table S5 for "Neurobehavioral effects of focused ultrasound-mediated blood-brain barrier opening"

**Supplementary Table S5. Additional pairwise comparisons for imaging and histology readouts in Fig. 4.** Results of 1-way ANOVAs with multiple comparisons corrections. * p<0.05; ** p<0.01; *** p<0.001; **** p<0.0001.

| Group1 | Group 2 | Summary (BBBO) | Adjusted P (BBBO) | Summary (SWI) | Adjusted P (SWI) | Summary (GFAP) | Adjusted P (GFAP) |
|---|---|---|---|---|---|---|---|
| **sham** | **High/D** | **** | <0.0001 | *** | 0.0002 | ** | 0.0018 |
| **sham** | **High/S** | **** | <0.0001 | ns | 0.9993 | * | 0.0138 |
| **sham** | **Ultra-High/D** | **** | <0.0001 | **** | <0.0001 | **** | <0.0001 |
| **sham** | **Medium/D** | **** | <0.0001 | ns | 0.9986 | ** | 0.0068 |
| **High/D** | **High/S** | ns | 0.9419 | **** | <0.0001 | ns | 0.9361 |
| **High/D** | **Ultra-High/D** | ns | 0.4865 | *** | 0.0005 | **** | <0.0001 |
| **High/D** | **Medium/D** | **** | <0.0001 | ** | 0.0004 | ns | >0.9999 |
| **High/S** | **Ultra-High/D** | ns | 0.1879 | **** | <0.0001 | **** | <0.0001 |
| **High/S** | **Medium/D** | **** | <0.0001 | ns | 0.9837 | ns | 0.9752 |
| **Ultra-High/D** | **Medium/D** | **** | <0.0001 | **** | <0.0001 | *** | 0.0002 |
