## Supplementary Table S6 for "Neurobehavioral effects of focused ultrasound-mediated blood-brain barrier opening"

**Supplementary Table S6. Additional comparisons for Supplementary Figure S4.**

| Groups | BBB opening (adjusted P values) | SWI (adjusted P values) | GFAP (adjusted P values) |
|---|---|---|---|
| CTX | | | |
| Medium/D vs. High/S | **<0.0001** | 0.9998 | 0.0901 |
| Medium/D vs. High/D | **<0.0001** | 0.063 | **0.0282** |
| Medium/D vs. Ultra-High/D | **<0.0001** | **0.0007** | 0.1853 |
| High/S vs. High/D | 0.9946 | 0.0518 | 0.9579 |
| High/S vs. Ultra-High/D | 0.9172 | **0.0005** | **<0.0001** |
| High/D vs. Ultra-High/D | 0.8091 | 0.3441 | **<0.0001** |
| DSC | | | |
| Medium/D vs. High/S | **0.0065** | >0.9999 | 0.5298 |
| Medium/D vs. High/D | **<0.0001** | **<0.0001** | **0.0219** |
| Medium/D vs. Ultra-High/D | **0.0314** | **<0.0001** | **<0.0001** |
| High/S vs. High/D | **<0.0001** | **<0.0001** | 0.3161 |
| High/S vs. Ultra-High/D | **<0.0001** | **<0.0001** | **<0.0001** |
| High/D vs. Ultra-High/D | **0.0135** | **0.0006** | **0.0069** |
| VSC | | | |
| Medium/D vs. High/S | 0.908 | >0.9999 | 0.9904 |
| Medium/D vs. High/D | **<0.0001** | 0.0526 | 0.5332 |
| Medium/D vs. Ultra-High/D | 0.3883 | **<0.0001** | **0.0038** |
| High/S vs. High/D | **<0.0001** | 0.051 | 0.6467 |
| High/S vs. Ultra-High/D | 0.1175 | **<0.0001** | **0.003** |
| High/D vs. Ultra-High/D | **<0.0001** | **0.0112** | 0.0911 |
| HC | | | |
| Medium/D vs. High/S | | | 0.7957 |
| Medium/D vs. High/D | | | 0.9294 |
| Medium/D vs. Ultra-High/D | | | **0.0007** |
| High/S vs. High/D | | | 0.9858 |
| High/S vs. Ultra-High/D | | | **<0.0001** |
| High/D vs. Ultra-High/D | | | **<0.0001** |
| Medium/D | | | |
| CTX vs. DSC | 0.872 | 0.8408 | **0.0004** |
| CTX vs. VSC | **0.0029** | 0.9519 | **0.0007** |
| DSC vs. VSC | **0.0123** | 0.9627 | 0.9989 |
| HC vs. CTX | | | 0.8456 |
| HC vs. DSC | | | 0.171 |
| HC vs. VSC | | | 0.2095 |
| High/S | | | |
| CTX vs. DSC | **<0.0001** | 0.8746 | 0.8364 |
| CTX vs. VSC | **<0.0001** | 0.9707 | 0.2606 |
| DSC vs. VSC | 0.9724 | 0.9631 | 0.752 |
| HC vs. CTX | | | 0.9783 |

| | | | |
|---|---|---|---|
| HC vs. DSC | | | 0.9975 |
| HC vs. VSC | | | 0.8065 |
| High/D | | | |
| CTX vs. DSC | 0.8959 | 0.0981 | 0.5092 |
| CTX vs. VSC | 0.8727 | 0.9667 | 0.9986 |
| DSC vs. VSC | 0.9986 | 0.0569 | 0.4121 |
| HC vs. CTX | | | 0.9812 |
| HC vs. DSC | | | 0.9319 |
| HC vs. VSC | | | 0.9606 |
| Ultra-High/D | | | |
| CTX vs. DSC | **0.0242** | **<0.0001** | 0.8836 |
| CTX vs. VSC | **<0.0001** | 0.4038 | **0.0086** |
| DSC vs. VSC | **0.0003** | **0.0048** | 0.0681 |
| HC vs. CTX | | | **0.0286** |
| HC vs. DSC | | | **0.0066** |
| HC vs. VSC | | | **<0.0001** |
