## Supplementary Table S7 for "Neurobehavioral effects of focused ultrasound-mediated blood-brain barrier opening"

**Supplementary Table S7*: Statistical Results for Figure 5.***

| Figure | Description | Samples | Test | Results |
|---|---|---|---|---|
| Fig. 5B | Comparison of ratios between post- and pre-treatment for fraction of trials close to target across treatments and days | Rat numbers:<br>Sham (S): 6<br>High/S (H/S): 6<br>High/D (H/D): 7<br>Ultra-High/D (UH): 7 | Linear Mixed Model: (Trials_close_to_target ~ treatment * days + (1\|Rat)); Estimated marginal means; mvt adjustment | Only contrasts with p<0.5 shown:<br>By Day (D):<br>**1) D1 S-H/D: p=0.0026**<br>**2) D1 S-UH: p=0.0003**<br>3) D1 H/D-H/S: p=0.3562<br>**4) D1 UH-H/S: p=0.043**<br>**5) D2 S-H/D: p=0.001**<br>6) D2 H/S-UH: p=0.1896<br>**7) D2 UH-H/S: p=0.0015**<br>**8) D3 S-UH: p=0.0009**<br>9) D3 H/D-UH: p=0.0988<br>**10) D3 UH-H/S: p=0.0002**<br>**11) D4 S-UH: p=0.001**<br>**12) D4 UH-H/S: p=0.0208**<br>13) D5 S-H/D: p=0.463<br>**14) D5 S-UH: p=0.0101**<br>15) D5 UH-H/S: p=0.1527<br>**16) D6 S-UH: p=0.0211**<br>17) D6 UH-H/S: p=0.2652<br>18) D7 S-UH: p=0.2661<br>19) D7 UH-H/S: p=0.1293<br>20) D8 S-UH: p=0.1451<br>**21) D9 S-UH: p=0.0241**<br>**22) D9 UH-H/S: p=0.0083**<br>23) D13 S-H/D: p=0.1064<br>24) D14 S-H/D: p=0.3903<br><br>By Treatment:<br>H/D<br>**1) D1-D2: p=0.0452**<br>**2) D1-D3: p=0.005**<br>**3) D1-D4: p=0.017**<br>**4) D1-D5: p=0.0224**<br>**5) D1-D6: p=0.011**<br>6) D1-D7: p=0.1837<br>**7) D1-D8: p=0.0172**<br>**8) D1-D9: p=0.0009**<br>**9) D1-D10: p=0.0001**<br>**10) D1-D11: p=0.0012**<br>**11) D1-D12: p=0.001**<br>12) D1-D13: p=0.2791<br>13) D1-D14: p=0.1963<br>**14) D1-D15: p=0.0005**<br>UH<br>1) D1-D7: p=0.2712 |

| | | | | |
|---|---|---|---|---|
| | | | | 2) D1-D8: p=0.1762<br>**3) D1-D10: p=0.0002**<br>**4) D1-D11: p<0.0001**<br>**5) D1-D12: p<0.0001**<br>**6) D1-D13: p<0.0001**<br>**7) D1-D14: p<0.0001**<br>**8) D1-D15: p<0.0001**<br>**9) D2-D10: p=0.0033**<br>**10) D2-D11: p=0.0005**<br>**11) D2-D12: p=0.0003**<br>**12) D2-D13: p<0.0001**<br>**13) D2-D14: p<0.0001**<br>**14) D2-D15: p<0.0001**<br>**15) D3-D10: p=0.0125**<br>**16) D3-D11: p=0.0024**<br>**17) D3-D12: p=0.0013**<br>**18) D3-D13: p=0.0001**<br>**19) D3-D14: p<0.0001**<br>**20) D3-D15: p=0.0003**<br>21) D4-D10: p=0.1572<br>**22) D4-D11: p=0.0409**<br>**23) D4-D12: p=0.0223**<br>**24) D4-D13: p=0.0025**<br>**25) D4-D14: p=0.0002**<br>**26) D4-D15: p=0.0056**<br>27) D5-D13: p=0.1521<br>**28) D5-D14: p=0.026**<br>29) D5-D15: p=0.3291<br>30) D6-D13: p=0.197<br>**31) D6-D14: p=0.0351**<br>32) D6-D15: p=0.4043<br>33) D7-D14: p=0.1581<br>34) D8-D14: p=0.2451<br>35) D9-D13: p=0.1376<br>**36) D9-D14: p=0.0212**<br>37) D9-D15: p=0.2774 |
| Fig. 5D | Comparison of number of missed sessions per night across treatments and days | Rat numbers: Sham (S): 6 High/S (H/S): 6 High/D (H/D): 7 Ultra-High/D (UH): 7 | Linear Mixed Model: (Missed_ sessions ~ treatment * days + (1\|Rat)); Estimated marginal means; mvt adjustment | Only contrasts with p<0.5 shown:<br>By Day (D):<br>1) D1 S-H/D: p=0.2103<br>**2) D1 S-UH: p=0.0008**<br>3) D1 H/S-H/D: p=0.2265<br>**4) D1 UH-H/S: p=0.0007**<br>**5) D2 S-UH: p=0.0004**<br>**6) D2 H/D-UH: p=0.0018**<br>**7) D2 UH-H/S: p=0.0441**<br>8) D3 S-UH: p=0.339<br>9) D3 H/D-UH: p=0.2644<br>10) D3 UH-H/S: p=0.3323 |

| | | | | |
|---|---|---|---|---|
| | | | | By Treatment:<br>H/D<br>**11) D1-D2: p=0.0181**<br>**12) D1-D3: p=0.0014**<br>**13) D1-D4: p=0.0016**<br>**14) D1-D5: p=0.0176**<br>**15) D1-D6: p=0.0014**<br>16) D1-D8: p=0.1369<br>**17) D1-D9: p=0.0015**<br>**18) D1-D10: p=0.0017**<br>**19) D1-D11: p=0.0015**<br>**20) D1-D12: p=0.0016**<br>**21) D1-D15: p=0.0015**<br>UH<br>1) D1-D3: p=0.4391<br>**2) D1-D4: p=0.0179**<br>**3) D1-D5: p<0.0001**<br>**4) D1-D6: p<0.0001**<br>**5) D1-D7: p=0.0004**<br>**6) D1-D8: p=0.0003**<br>**7) D1-D9: p<0.0001**<br>**8) D1-D10: p<0.0001**<br>**9) D1-D11: p<0.0001**<br>**10) D1-D12: p<0.0001**<br>**11) D1-D13: p<0.0001**<br>**12) D1-D14: p<0.0001**<br>**13) D1-D15: p<0.0001**<br>**14) D2-D5: p=0.0029**<br>**15) D2-D6: p=0.0029**<br>**16) D2-D7: p=0.0402**<br>**17) D2-D8: p=0.0382**<br>**18) D2-D9: p=0.0013**<br>**19) D2-D10: p=0.0001**<br>**20) D2-D11: p=0.0001**<br>**21) D2-D12: p=0.0029**<br>**22) D2-D13: p=0.0001**<br>**23) D2-D14: p=0.0002**<br>**24) D2-D15: p=0.0015**<br>**25) D3-D10: p=0.3451**<br>**26) D3-D11: p=0.3447**<br>**27) D3-D13: p=0.3448**<br>**28) D3-D14: p=0.3443** |
