## Supplementary Table S8 for "Neurobehavioral effects of focused ultrasound-mediated blood-brain barrier opening"

**Supplementary Table S8*:* Statistical Results for Supplementary Figure S5.**

| Figure | Description | Samples | Test | Results |
|---|---|---|---|---|
| Fig. S5A | Comparison of ratios between post- and pre-treatment for IPIs across treatments and days | Rat numbers:<br>Sham (S): 6<br>High/S (H/S): 6<br>High/D (H/D): 7<br>Ultra-High/D (UH): 7 | Linear Mixed Model:<br>(IPI ~ treatment * days + (1\|Rat));<br>Estimated marginal means; mvt adjustment | Only contrasts with p<0.5 shown:<br>By Day (D):<br>**1) D1 S-H/D: p=0.0173**<br>**2) D2 S-UH: p=0.0045**<br>**3) D2 H/S-UH: p=0.0098**<br>**4) D3 S-UH: p=0.0373**<br>**5) D3 UH-H/S: p=0.0016**<br>6) D4 UH-H/S: p=0.2052<br>7) D5 UH-H/S: p=0.2245<br>8) D6 UH-H/D: p=0.1981<br>**9) D6 UH-H/S: p=0.0293**<br>10) D7 UH-H/S: p=0.4208<br>11) D8 UH-H/S: p=0.3275<br>12) D13 UH-H/S: p=0.3163<br><br>By Treatment:<br>H/D<br>1) D1-D5: p=0.4767<br>**2) D1-D6: p=0.0023**<br>3) D1-D7: p=0.0754<br>4) D1-D8: p=0.0882<br>5) D1-D10: p=0.085<br>**6) D1-D11: p=0.0476**<br>7) D1-D12: p=0.1012<br>8) D1-D13: p=0.2303<br>9) D1-D14: p=0.3947<br>**10) D1-D15: p=0.0303**<br>11) D2-D6: p=0.2137<br>12) D3-D6: p=0.2298<br>UH<br>1) D2-D9: p=0.1928<br>2) D2-D10: p=0.2817<br>3) D2-D12: p=0.3916<br>4) D2-D13: p=0.3398<br>5) D2-D15: p=0.2037<br>6) D3-D9: p=0.3953<br>7) D3-D10: p=0.4755<br>8) D3-D15: p=0.4136 |
| Fig. S5B | Comparison of ratios between post- and pre-treatment for CVs of IPIs across treatments and days | Rat numbers:<br>Sham (S): 6<br>High/S (H/S): 6<br>High/D (H/D): 7<br>Ultra-High/D (UH): 7 | Linear Mixed Model:<br>(CV ~ treatment * days + (1\|Rat)); | Only contrasts with p<0.5 shown:<br>By Day (D):<br>**1) D1 S-H/D: p<0.0001**<br>2) D1 S-H/S: p=0.256<br>3) D1 H/S-H/D: p=0.3661<br>**4) D3 S-UH: p=0.0166**<br>5) D3 H/D-UH: p=0.4738 |

| | | | | |
|---|---|---|---|---|
| | | | Estimated marginal means; mvt adjustment | **6) D3 H/S-UH: p=0.0004**<br>**7) D4 S-UH: p=0.02**<br>8) D4 H/D-UH: p=0.3707<br>**9) D4 H/S-UH: p=0.001**<br>**10) D5 S-UH: p=0.0012**<br>11) D5 H/D-UH: p=439<br>**12) D5 H/S-UH: p<0.0001**<br>**13) D6 S-UH: p=0.0041**<br>14) D6 H/D-UH: p=0.1756<br>**15) D6 H/S-UH: p=0.0003**<br>16) D7 S-UH: p=0.4306<br>17) D7 H/S-UH: p=0.1138<br>18) D8 H/S-UH: p=0.195<br>19) D9 S-UH: p=0.0787<br>20) D9 H/D-UH: p=0.4039<br>**21) D9 H/S-UH: p=0.0019**<br>22) D10 H/S-UH: p=0.0569<br>**23) D11 S-UH: p=0.0355**<br>**24) D11 H/S-UH: p=0.0096**<br><br>By Treatment:<br>H/D<br>**25) D1-D2: p=0.0008**<br>**26) D1-D3: p=0.0006**<br>**27) D1-D4: p=0.0001**<br>**28) D1-D5: p=0.002**<br>**29) D1-D6: p<0.0001**<br>**30) D1-D7: p=0.004**<br>**31) D1-D8: p<0.0001**<br>**32) D1-D9: p<0.0001**<br>**33) D1-D10: p<0.0001**<br>**34) D1-D11: p<0.0001**<br>**35) D1-D12: p<0.0001**<br>**36) D1-D13: p=0.0002**<br>**37) D1-D14: p<0.0001**<br>**38) D1-D15: p<0.0001**<br>UH<br>13) D3-D12: p=0.4832<br>14) D3-D13: p=0.4681<br>15) D3-D15: p=0.3430<br>16) D4-D15: p=0.4622<br>17) D5-D12: p=0.1765<br>18) D5-D13: p=0.1674<br>19) D5-D14: p=0.2320<br>20) D5-D15: p=0.0947<br>21) D6-D12: p=0.455<br>22) D6-D13: p=0.4386<br>23) D6-D15: p=0.2954 |

| | | | | |
|---|---|---|---|---|
| | | | | H/S<br>1) D1-D2: p=0.0985<br>**2) D1-D3: p=0.0164**<br>**3) D1-D4: p=0.0229**<br>**4) D1-D5: p=0.0133**<br>**5) D1-D6: p=0.0192**<br>**6) D1-D7: p=0.0161**<br>**7) D1-D8: p=0.0083**<br>**8) D1-D9: p=0.0072**<br>**9) D1-D10: p=0.0047**<br>**10) D1-D11: p=0.0027**<br>**11) D1-D12: p=0.0171**<br>**12) D1-D13: p=0.007**<br>13) D1-D14: p=0.0647<br>**14) D1-D15: p=0.0127** |
