## Supplementary Table S9 for "Neurobehavioral effects of focused ultrasound-mediated blood-brain barrier opening"

**Supplementary Table S9*: Statistical Results for Supplementary Figure S6*.**

| Figure | Description | Samples | Test | Results |
|---|---|---|---|---|
| Fig. S6A | Comparison of ratios between post- and pre-treatment for fraction of trials close to target across treatments and days | Rat numbers:<br>Sham (S): 6<br>Medium/S (M/S): 5<br>Medium/I (M/I): 5<br>Medium/D (M/D): 5 | Linear Mixed Model: (Trials_close_to_target ~ treatment * days + (1\|Rat)); Estimated marginal means; mvt adjustment | Only contrasts with p<0.5 shown:<br>By Day (D):<br>1) D1 M/D-M/I: p=0.3211<br>**2) D1 M/I-S: p=0.0231**<br>**3) D1 M/S-S: p=0.001**<br>4) D15 M/D-M/S: p=0.3709<br>By Treatment:<br>M/I<br>**1) D1-D2: p=0.0062**<br>2) D1-D3: p=0.2652<br>**3) D1-D4: p=0.0907**<br>**4) D1-D5: p=0.0175**<br>**5) D1-D6: p=0.0128**<br>**6) D1-D7: p=0.0819**<br>7) D1-D11: p=0.1718<br>M/S<br>1) D1-D2: p=0.1445<br>**2) D1-D3: p=0.0002**<br>**3) D1-D4: p<0.0001**<br>**4) D1-D5: p=0.0002**<br>**5) D1-D6: p=0.0001**<br>**6) D1-D7: p<0.0001**<br>**7) D1-D8: p<0.0001**<br>**8) D1-D9: p<0.0001**<br>**9) D1-D10: p<0.0001**<br>**10) D1-D11: p<0.0001**<br>**11) D1-D12: p=0.0282**<br>**12) D1-D13: p=0.0348**<br>**13) D1-D14: p=0.0007**<br>**14) D1-D15: p<0.0001** |
| Fig. S6B | Comparison of number of missed sessions per night across treatments and days | Rat numbers:<br>Sham (S): 6<br>Medium/S (M/S): 5<br>Medium/I (M/I): 5<br>Medium/D (M/D): 5 | Linear Mixed Model: (Missed_sessions ~ treatment * days + (1\|Rat)); Estimated marginal means; mvt adjustment | Only contrasts with p<0.5 shown:<br>By Day (D):<br>1) D1 M/S-S: p=0.0594<br><br>By Treatment:<br>M/S<br>**2) D1-D2: p=0.0407**<br>**3) D1-D3: p=0.001**<br>**4) D1-D4: p=0.0016**<br>**5) D1-D5: p=0.0017**<br>**6) D1-D6: p=0.0015**<br>**7) D1-D7: p=0.0017**<br>**8) D1-D8: p=0.0017**<br>**9) D1-D9: p=0.0017** |

| | | | | |
|---|---|---|---|---|
| | | | | **10) D1-D10: p=0.0017**<br>**11) D1-D11: p=0.0018**<br>12) D1-D12: p=0.3822<br>13) D1-D13: p=0.382<br>**14) D1-D14: p=0.0018**<br>**15) D1-D15: p=0.0018** |
| Fig. S6C | Comparison of ratios between post- and pre-treatment for IPIs across treatments and days | Rat numbers:<br>Sham (S): 6<br>Medium/S (M/S): 5<br>Medium/I (M/I): 5<br>Medium/D (M/D): 5 | Linear Mixed Model:<br>(IPI ~ treatment * days + (1\|Rat));<br>Estimated marginal means; mvt adjustment | Only contrasts with p<0.5 shown:<br>By Day (D):<br>1) D1 M/D-M/I: p=0.1595<br>**2) D2 M/S-S: p=0.0354**<br><br>By Treatment:<br>M/I<br>8) D1-D15: p=0.4053<br>M/S<br>**15) D1-D2: p=0.0307**<br>16) D2-D9: p=0.4774<br>17) D2-D10: p=0.3626<br>18) D2-D14: p=0.2283<br>S<br>1) D1-D9: p=0.2356 |
| Fig. S6D | Comparison of ratios between post- and pre-treatment for CVs of IPIs across treatments and days | Rat numbers:<br>Sham (S): 6<br>Medium/S (M/S): 5<br>Medium/I (M/I): 5<br>Medium/D (M/D): 5 | Linear Mixed Model:<br>(CV ~ treatment * days + (1\|Rat));<br>Estimated marginal means; mvt adjustment | Only contrasts with p<0.5 shown:<br>By Day (D):<br>16) D1 M/S-S: p=0.1559<br>17) D14 M/D-M/S: p=0.3951<br>**18) D14 M/D-S: p=0.0109**<br><br>By Treatment:<br>M/D<br>19) D6-D14: p=0.347<br>20) D7-D14: p=0.3488<br>21) D13-D14: p=0.2877<br>M/S<br>3) D1-D4: p=0.294<br>4) D1-D7: p=0.1012<br>**5) D1-D8: p=0.0455**<br>6) D1-D9: p=0.0916<br>7) D1-D10: p=0.0702<br>8) D1-D11: p=0.1842<br>9) D1-D12: p=0.3147<br>10) D1-D13: p=0.4303<br>11) D1-D14: p=0.4146<br>12) D1-D15: p=0.2603 |
