## Supplementary Table S10 for "Neurobehavioral effects of focused ultrasound-mediated blood-brain barrier opening"

**Supplementary Table S10*: Statistical Results for Figure 6.***

| Figure | Description | Samples | Test | Results |
|---|---|---|---|---|
| Fig. 6B | Comparison of Trial-by-Trial correlations for movement trajectories before MB-FUS treatment, across treatment groups and across timepoints (before treatment, Day 1 after treatment, Day 15 after treatment). | Rat numbers:<br>Sham (S): 6<br>High/S (H/S): 6<br>High/D (H/D): 7<br>Ultra-High/D (UH): 7 | Linear Mixed Model: (Trial-by-trial correlations ~ treatment * days + (1\|Rat)); Estimated marginal means; mvt adjustment | Only contrasts with p<0.5 shown:<br>By Day (D):<br>1) D1 H/S-H/D: p=0.1909<br>**2) D1 S-H/D: p=0.0461**<br>**3) D1 H/S-UH: p=0.0076**<br>**4) D1 S-UH: p=0.00008**<br><br>By Treatment:<br>**1) H/D Bef-D1: p=0.0336**<br>2) H/D D1-D15: p=0.1035<br>**3) UH Bef-D1: p=0.0001**<br>**4) UH D1-D15: p=0.0019** |
| Fig. 6C | Comparison of correlations between movement trajectories before treatment and on Day 1 or Day 15 after treatment across groups and days. Correlations are normalized to the pre-treatment trial-by-trial correlations (shown in Fig. 5B). | Rat numbers:<br>Sham (S): 6<br>High/S (H/S): 6<br>High/D (H/D): 7<br>Ultra-High/D (UH): 7 | Linear Mixed Model: (Normalized Pre-Post Correlations ~ treatment * days + (1\|Rat)); Estimated marginal means; mvt adjustment | Only contrasts with p<0.5 shown:<br>By Day (D):<br>1) D1 S-H/D: p=0.489<br>**2) D1 S-UH: p=0.0039**<br>3) D1 H/D-UH: p=0.2475<br>4) D1 S-H/S: p=0.1658<br>5) D15 S-H/S: p=0.34<br><br>By Treatment:<br>**6) UH D1-D15: p=0.0048** |
