## Supplementary Table S11 for "Neurobehavioral effects of focused ultrasound-mediated blood-brain barrier opening"

**Supplementary Table S11*: Statistical Results for Supplementary Figure S8.***

| Figure | Description | Samples | Test | Results |
|---|---|---|---|---|
| Fig. S8A | Comparison of Trial-by-Trial correlations for movement trajectories before MB-FUS treatment, across treatment groups and across timepoints (before treatment, Day 1 after treatment, Day 15 after treatment). | Rat numbers: Sham (S): 6 Medium/S (M/S): 5 Medium/I (M/I): 5 Medium/D (M/D): 5 | Linear Mixed Model: (Trial-by-trial correlations ~ treatment * days + (1\|Rat)); Estimated marginal means; mvt adjustment | Only contrasts with p<0.5 shown:<br>By Day (D):<br>1) D1 S-M/S: p=0.0869<br>2) D1 M/S-M/I: p=0.3723<br>3) D15 S-M/S: p=0.2796<br><br>By Treatment:<br>**1) M/S Bef-D1: p=0.0339**<br>2) M/S D1-D15: p=0.3424 |
| Fig. S8B | Comparison of correlations between movement trajectories before treatment and on Day 1 or Day 15 after treatment across groups and days. Correlations are normalized to the pre-treatment trial-by-trial correlations (shown in Fig. S8A). | Rat numbers: Sham (S): 6 Medium/S (M/S): 5 Medium/I (M/I): 5 Medium/D (M/D): 5 | Linear Mixed Model: (Normalized Pre-Post Correlations ~ treatment * days + (1\|Rat)); Estimated marginal means; mvt adjustment | Only contrasts with p<0.5 shown:<br>By Day (D):<br>1) D1 S-M/S: p=0.0942<br>2) D1 M/S-M/I: p=0.2578<br>3) D1 M/S-M/D: p=0.2742<br><br>By Treatment:<br>4) M/S D1-D15: p=0.2622 |
