## Supplementary Table S12 for "Neurobehavioral effects of focused ultrasound-mediated blood-brain barrier opening"

**Supplementary Table S12. Additional statistical results for Figure 7.**

| Group1 | Group 2 | Summary (SWI) | Adjusted P value (SWI) | Summary (T2w) | Adjusted P value (T2w) |
|---|---|---|---|---|---|
| Day 0 | Day 1 | **** | <0.0001 | **** | <0.0001 |
| Day 0 | Day 3 | **** | <0.0001 | **** | <0.0001 |
| Day 0 | Day 7 | **** | <0.0001 | **** | <0.0001 |
| Day 0 | Day 14 | ** | 0.0011 | *** | 0.0001 |
| Day 0 | Day 21 | ** | 0.0010 | **** | <0.0001 |
| Day 1 | Day 3 | ns | 0.9937 | ns | >0.9999 |
| Day 1 | Day 7 | ns | 0.1153 | ** | 0.0013 |
| Day 1 | Day 14 | ** | 0.0031 | ns | 0.9475 |
| Day 1 | Day 21 | * | 0.0478 | ns | >0.9999 |
| Day 3 | Day 7 | ** | 0.0056 | * | 0.0137 |
| Day 3 | Day 14 | **** | <0.0001 | ns | 0.9628 |
| Day 3 | Day 21 | ** | 0.0014 | ns | >0.9999 |
| Day 7 | Day 14 | ns | 0.1789 | ** | 0.0056 |
| Day 7 | Day 21 | ns | 0.4426 | * | 0.0144 |
| Day 14 | Day 21 | ns | 0.9993 | ns | 0.8271 |
